# Open Hardware for neuro-prosthesis research: A study about a closed-loop multi-channel system for electrical surface stimulations and measurements

**DOI:** 10.1101/141184

**Authors:** David Rotermund, Udo A. Ernst, Klaus R. Pawelzik

## Abstract

Recent progress in neuro-prosthetic technology gives rise to the hope that in the future blind people might regain some degree of visual perception. It was shown that electrically stimulating the brain can be used to produce simple visual impressions of light blobs (phosphenes). However, this perception is very far away from natural sight. For developing the next generation of visual prostheses, real-time closed-loop stimulators which measure the actual neuronal activities and on this basis determine the required stimulation pattern. This leads to the challenge to design a system that can produce arbitrary stimulation-patterns with up to ±70V and with up to 25mA while measuring neuronal signals with amplitudes in the order of mV. Furthermore, the interruption of the measurement by stimulation must be as short as possible and the system needs to scale to hundreds of electrodes. We discuss how such a system and especially its current pumps and input protection need to be designed and which problems arise. We condense our findings into an example design for which we provide all design files (boards, firmwares and software) as open-source. This is a first step in taking the existing open-source www.open-ephys.org recording system and converting it into a closed-loop experimental setup for neuro-prosthetic research.

## 2 Introduction

Loosing the ability to see is a traumatic experience that can happen to everyone [1]. Using technology replacing lost senses, like touch (e.g. [2, 3, 4, 5]), hearing (e.g. [6, 7, 8, 9]) and vision (e.g. [10, 11, 12, 13, 14, 15, 16]), is a goal that is pursued by many researchers since many decades.

In the field of vision restoration, which we are especially interested in, two main research directions can be observed: The first one is covered by retinal implants [12, 17, 18, 19, 20, 21, 22, 23] which use still functional sub-layers of the patient’s retina. These kind of implants show very promising results in first clinical trials and allow suitable patients to interact with their environment, based on the information delivered to the remains of the retina [24, 25]. However, the requirement on the patient’s eyes is not met by many of the blind patients [26]. This makes the second approach, which is based on the idea of a visual cortex prostheses [27, 28, 29, 30], very appealing. Here, the visual information is sent directly into the brain by means of electrical stimulation of visual cortex’s nerve cells. In this paper we will focus on these visual cortex prostheses, but the presented technology can be applied to research on prostheses for the other senses, too.

The idea behind a typical sensory cortex prosthesis is simple [11]: Record the physical signal via a measurement device (e.g. camera, microphone or touch sensor) and digitize it. Based on the recorded time series, calculate a suitable stimulation pattern in real-time. Convert these patterns into electrical stimulation currents [31] via current pumps [32, 33] or, in the case of opto-genetic stimulation [34, 7], into a light signal which is applied to genetically modified neuronal cells. For normal electrical stimulation, which will be discussed here, the electrical currents are conducted to electrodes (e.g. [35, 36, 37, 38, 39, 40, 41, 42, 43]) which interface the brain tissue [44]. Finally, the current reaches the nerve cells. This external intervention changes the activity of the ongoing neuronal activities and should, if done correctly, result in the desired perception.

In reality, the realization of this idea is not that simple. Alone the question, how and where to stimulate is still a field of research (e.g. [45, 46, 47, 48, 49, 50, 51, 52, 53, 54, 55, 56]), with the goal to produce brain activity patterns similar to those evoked by sensory stimulation. Furthermore, the activity patterns produced by electric stimulation should be processed by the brain similar to natural brain activity and should result in natural perceptions. This is a largely unsolved problem and it is not clear if this goal can ever be fully reached [45]. Nevertheless, in the field of electrical stimulation-based hearing aids the actual state of stimulation is very promising. Current development of cortical vision prostheses did not went beyond simple light blobs (phosphenes) [47, 49]. Our long term research agenda is to develop a technological and scientific foundation for improving the situation. One idea is to use stimulation in a closed-loop configuration [57], possibly even with under-threshold stimulation [58, 59, 60], as well as to exploit the brain’s capability to adapt [61, 62].

For a closed-loop approach, the proposed technology (see figure 1) is composed of three parts: 1.) Measuring neuronal activity patterns, which requires bio-signal amplifiers [63, 64, 65] and analog-digital converters suitable for neuronal signals [66, 67, 68, 69]. 2.) A data processor [70] for evaluating the actual state of the neuronal network as well as for predicting the required intervention to bring the neuronal network into the desired dynamical state. 3.) The electrical stimulator itself [68, 71, 33]. All three parts of the system must work and interact in real-time.

**Figure 1:**
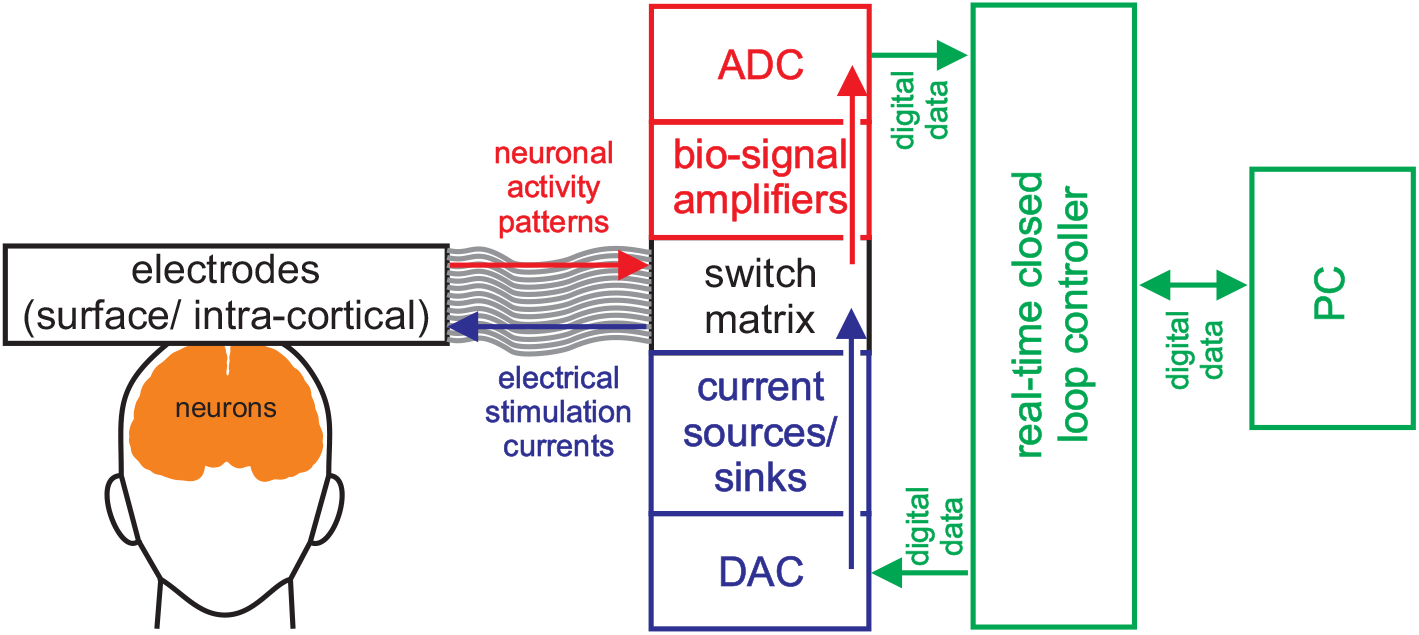
Sketch of a closed-loop system for neuro-prosthesis applications. Electrodes establish an interface to the brain tissue. On the measuring side of the system, the electrodes are connected to bio-amplifiers for boosting the signal strength of the observed neuronal mV-sized signals. These amplifiers send their amplified signals to an analog digital converter (ADC). The digitized stream of measurement data is conveyed to a real time closed loop controller. This controller analyses the incoming data and calculates which intervention is required. The results of these calculations are a spatial temporal stimulation pattern. This pattern is send to the digital analog converter (DAC) which controls current pumps that are connected to the electrodes which closes the loop.

The final goal would clearly be to build a wireless implantable system [72, 73, 74]. However, such an implant needs to fulfill a large number of requirements and needs to possess special properties (see discussion for details). This makes it much more efficient to start with an external closed-loop system for performing research on more advanced stimulation paradigms. Later, when it will be much clearer how a more natural stimulation needs to be done, the system can be miniaturize towards an implant.

Concerning visual cortex prostheses, two types of neuro-interfaces have been used in research [29]: Surface electrodes arrays [30, 75] and intra-cortical electrodes [76]. The presented system is optimized for surface electrodes but can be adapted for using intra-cortical electrodes. Specifically, with our system we aim for very high density and small size surface electrodes [42] with several hundreds of electrodes. For traditional surface electrode arrays used in a medical applications, the electrodes have a diameter up to 1cm. Stimulating the brain with such electrodes already requires voltages up to 20V [30] for delivering currents of several mA. For increasing the density of the phosphenes, the electrode diameter was reduced to 560*μ*m [42]. Some researchers involved in the development of these electrodes even expect that voltages up to 100V with currents of 15mA (or maybe even more) [33, 77] might be necessary. However, the optimal voltage and current range is not determined yet and may require lower voltages. We designed our system for voltages in the range of ± 70V and currents up to 25mA. Traditionally, simple mono- or bi-phasic stimulation pulse shapes are used [78]. Since the properties of the phosphenes might depend on the pulse shape, we designed our system to instead produce arbitrary pulse shapes with a time resolution of 40k samples/s.

To date, suitable systems with the described specifications are not commercially available. Only few systems are capable of producing high voltages and high currents or scale up to several hundreds of electrodes. Furthermore, these systems are closed source or are just not designed for a closed-loop application. Such a system needs to e.g. minimization of the duration between the last stimulation pulse and continuing the recording of the neuronal activities, analyzing the data in real-time, or continuously changing the stimulation patterns in real-time. As a result, embedding these systems into a real-time, low latency closed-loop system is a problem.

In the end, we decided to build our own system. However, we wanted to use the open-source recording system open-ephys (www.open-ephys.org) with its Intan RHD ICs (includes bio-amplifier and an ADC for neuronal signal recording applications; www.intantech.com) for the recording part. In this paper we describe our first step towards this goal and make all the corresponding design files available as open-source in the supplement materials.

## 3 Results

In the results section we will explain the ideas which went into the design of the system concept (Figure 2), problems that occurred and how we solved them. We also present an example design for the circuit diagrams as well as circuit board designs. The full system is partitioned into distinct modules that serve different functions: First we will describe the modules containing the current pumps which provide the stimulation current. In a second step, we combine these modules and add control structures for obtaining a 16-channel building block. In the next step, cascades of protection switches are added that will allow the combination of stimulation and measurement equipment into one system. Finally, we explain the structure of the necessary firmware(s) for controlling the system. The section ends with a summary of our experiences from tests of the system’s concept in real hardware. All design files of the system, its firmwares and software, as well as of the used test modules are part of the supplemental material.

**Figure 2:**
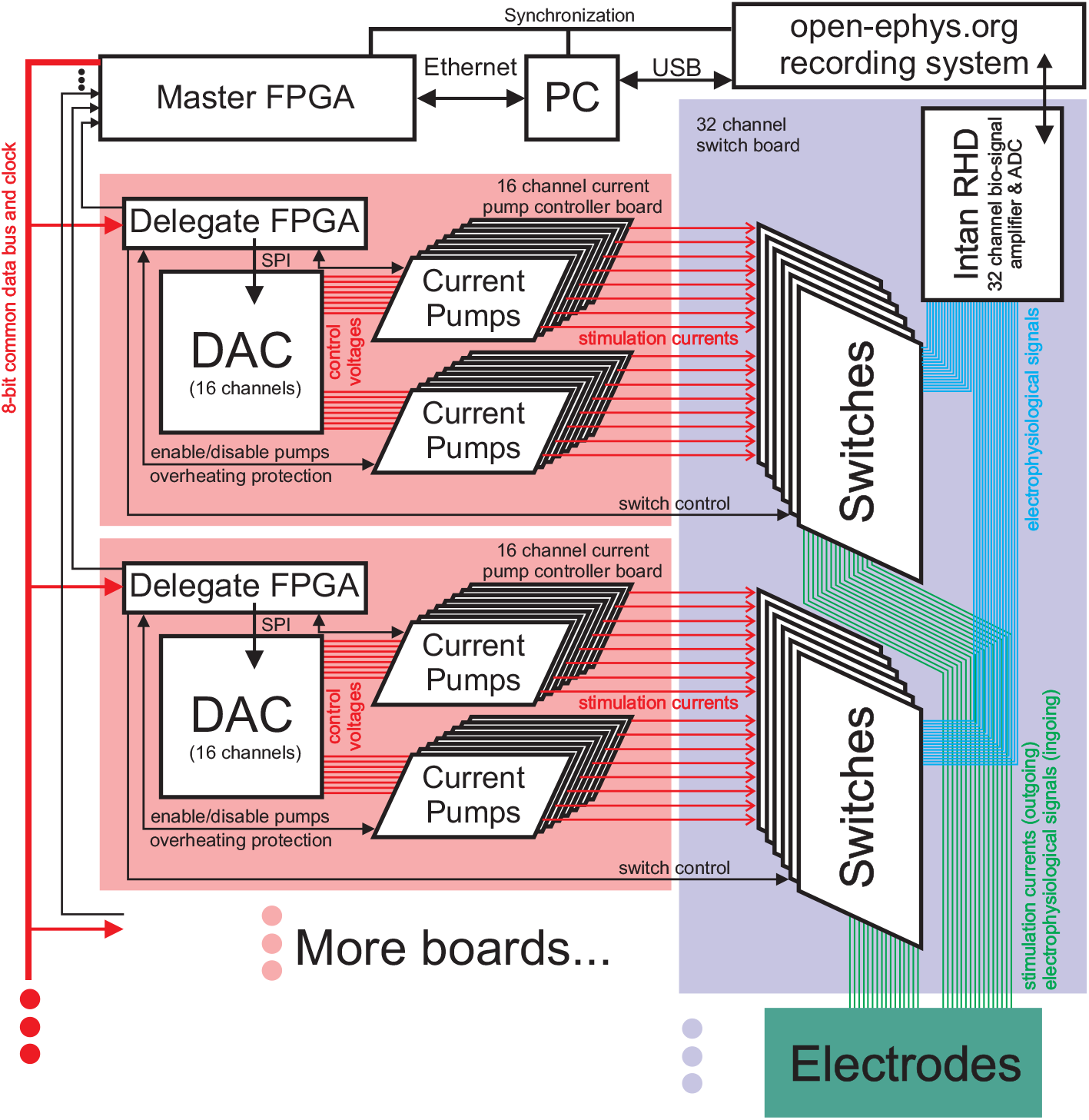
The system concept shows how the different building blocks (PC, Master FPGA, Open-Ephys recording system, 16 channel current pump controller boards with their submodules as well as the 32 channel switch boards) interact with each other.

### 3.1 Current pumps

Core of the stimulation system are the current sources which are realized as modules each containing a single current pump for easy replacement. For covering the envisioned applications, the current pump will deliver up to ±25mA using voltages up to ±70V.

#### Circuit design

As the basis for the stimulation system, we decided to use an ‘improved’ Howland current pump [79]. Figure 3(a) shows the circuit diagram of such a constant current pump. The circuit is controlled by an external DAC (Analog AD5360) providing an input voltage *V_In_* proportional to the desired output current. According to [79] the selection of the resistors needs to obey the equation

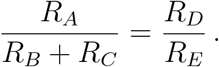

**Figure 3:**
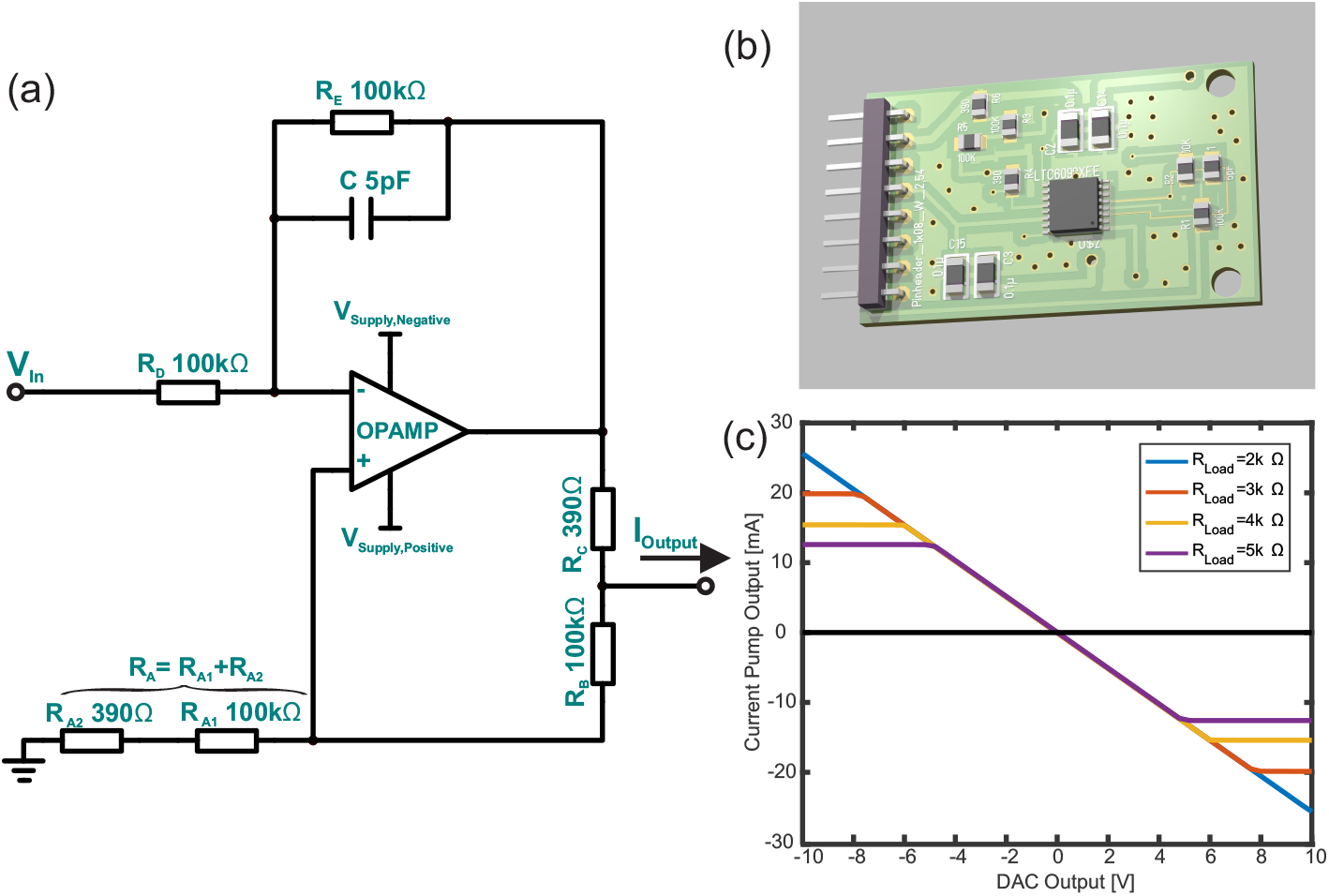
(a) Circuit diagram for an ‘improved’ Howland current pump [79]. Components are selected for an op-amp LTC6090 with ±70V as supply voltages and *V_In_* ∈ [−10*V*,…, 10*V*] as well as a maximum output current of ±25.6mA. (b) 3D rendering of an example design (double layer board with single side component load) of this circuit diagram (see supplement for the design files). (c) Simulation of the circuit diagram with ±70V as supply voltages in Texas Instruments TINA software. The graph shows the theoretical output current of the current pump in dependence of *V_In_* and for different resistive loads.

The application information in the datasheet for the selected op-amp LTC6090 suggests to use large feedback resistors. Following the manuals’ line of argument, a resistor with a value of 100kΩ was selected for *R_E_*. *R_D_* was also set to 100kΩ since the manual for the Analog AD5630 DAC suggests a load with a minimum of 10kΩ for the intended parameters of operation. Like in the example in [79], we set *R_A_*_1_ and *R_C_* to 100kΩ as well. The selection of the values for *R_C_*, as well as *R*_*A*2_, depends on the required maximal current and the maximal control voltage *V_In_* according to the equation

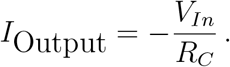

We expect the external DAC to deliver control voltages in the range of ±10V (with 16-bit resolution) and aimed for a maximal current of ≈ ±25mA, which is half of the output capabilities of a LTC6090 op-amp. Based on the available E12 values for resistors, we selected *R_C_* = *R*_*A*2_ = 390Ω thus setting the maximum current to ±25.6mA. Applying the maximum symmetric supply voltages of ±70V to the op-amp, we simulated the behavior of this current pump with different loads in Texas Instruments’ TINA software. Figure 3(c) displays the results of this simulation. Given the maximum driving voltage of ±70V, already at a resistive load of 3kΩ the maximum current output of the current pump is limited to a value lower than ±25.6mA. Increasing the resistive load further reduces the current, that can be maximally sourced by the current pump with the given supply voltage.

In figure 4 simulations are shown for two additional cases where the supply voltage was limited to more typical ±15V and the resistors *R_C_* = *R*_*A*2_ were selected such that the maximum currents were limited to 1mA and 100*μ*A. In the case of *I*_Output,Max_ = 100*μ*A, the current pump can drive resistive loads with 30kΩ. For these three examples, table 1 lists the necessary resistor values and the implied size of the addressable current steps via a 16-bit DAC. In applications where no high voltages (<35V) are required (e.g., for intra-cortical stimulation), the noise performance may be improved if the LTC6090 is replaced by a low noise, non high-voltage op-amp. However, the requirement for an input pin that can enable/disable the output and an indicator for overheating reduces the available op-amp IC types strongly.

**Figure 4:**
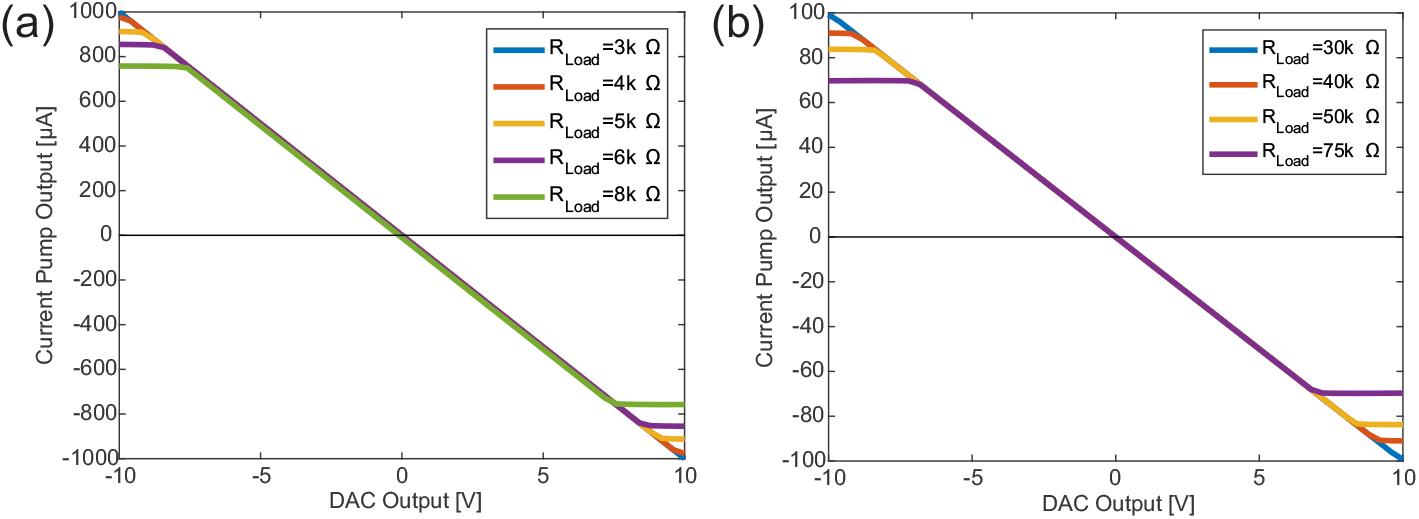
The graph shows the theoretical output current of the current pump in dependence of *V_In_* and for different resistive loads (see figure 3(c) for details). Here the circuit was simulated for supply voltages of ±15V and designed to produce a maximum current of 1mA (a) and 100*μ*A (b). The latter is in the order of magnitude of currents typically used for intra-cortical stimulations. Table 1 lists the configurations and properties of these circuits.

**Table 1:** Summarizing the results from the simulations shown in figure 3 and 4 as well as listing the used parameters.

| $R_C = R_{A2}$ | $V_{Supply}$ | $I_{Output,Max}$ | $\Delta I_{DAC\ step}$ |
| --- | --- | --- | --- |
| 390Ω | $\pm 70V$ | $\pm 25.6mA$ | $\approx 0.8\mu A$ |
| 10kΩ | $\pm 15V$ | $\pm 1mA$ | $\approx 0.03\mu A$ |
| 100kΩ | $\pm 15V$ | $\pm 100\mu A$ | $\approx 0.003\mu A$ |

#### Circuit calibration

It is necessary to calibrate the *I*_Output_(*V_In_*) curve for every individual channel, since production tolerances in the resistors cause significant differences to the theoretically expected behavior. This may lead to deviations in the shape of the *I*-*V_In_*-curves as well as to non-zero current flows at *V_In_* = 0. Software based calibration (i.e., re-mapping of the DAC input values to measured output currents) can help to compensate for aberrations of the curve’s shape down to Δ*I*_DAC step_ level. However, this might not be good enough for compensating the current flow at *V_In_* = 0. Even a small current can cause damage to the brain tissue and electrodes through electrophoresis and electrolysis, if it is applied for an extended time. As an easy solution, only op-amps with an enable/disable pin for the output should be used, hence allowing to turn off the current pump when it is not needed. Furthermore, it is highly advisable to place an additional analog switch in the output path of the current pump to prevent external currents from flowing out of or into the current pump during the time when the current source is disabled (see section ‘Analog protection switches’ for details).

If the current pump will be used with supply voltages of ±70V, special care is needed to keep electrical conductors sufficiently separated. According to IPC’s standard IPC-2221A [80], section 6.3 ‘Electrical Clearance’, their recommendation for an appropriate distance can be taken from table 6–1 or calculated (for larger voltages) with the equation

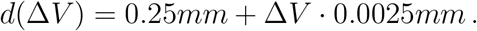

With Δ*V* = 140*V* we need at least 0.6mm, while the IPC table 6–1 even suggests 0.8mm (for up to Δ*V* = 250*V*). Hence for our designs presented in the supplement we used 0.75mm as electrical conductor spacing. For Δ*V* = 30*V* a spacing of only 0.25mm would be required. Using conformal coating can reduce the conductor spacing in both cases by half.

#### Module design

In figure 3(b) our example design is shown. A module contains one current pump on a two-layer printed circuit board. The bottom side is not loaded with components. Due to the required large conductor spacing as well as the larger resistor/capacitor package sizes which come with the higher voltage rating, the board has a size of 38mm x 23mm. For smaller Δ*V*, the necessary area for one current pump can be reduced significantly, also by using connectors with a smaller pitch.

Several of these modules can share four connections for supply voltages (positive and negative), digital ground and analog ground. The two inputs for *V_In_* and the enable/disable signal for the pump can not be shared since they are controlled for each current pump individually. Furthermore, two individual outputs are provided by every module, namely the stimulation current itself and a signal of the op-amp indicating overheating. Thus the number of connections *N*_Con_ for *N*_Channels_ channels scales as *N*_Con_ = 4 + 4 · *N*_Channels_.

In our trial production of test boards, we found one of three modules being non-functional. The problem could be traced back to the op-amp which showed erratic behavior. Due to the (large) contact pad on the flip side of the IC’s package and the huge copper areas for cooling the IC, it is very problematic to replace an individual op-amp especially if it is surrounded by other components. As as consequence, we suggest to put only a few of these current pumps with only the necessary support components on one board, thus allowing easy replacement by exchanging the whole board. For the design presented in the supplement, we decided to put only one current pump on a board to keep the financial damage low while trading it in for a larger required spatial volume.

### 3.2 Combining and controlling a multitude of current pumps

For an application like a visual prostheses, a system is required which can serve several hundreds of electrodes. The aim is to design elementary building blocks that can flexibly be combined to create systems ranging from a few up to a very large number (e.g. 256) of stimulation channels. In the following, we will present the design of a 16-channel building block, which consists of an FPGA (for a better discrimination between the different FPGAs, we will call it ‘delegate FPGA’) controlling a 16-channel digital-analog converter (DAC), which in turn delivers the output for controlling up to 16 current pumps of the type described in the last section. In parallel, this delegate FPGA is also handling the temperature warning signals of its current pumps, and enables/disables these current pumps as needed.

The design is based around the Analog Devices AD5360 which is a 16-channel digital-analog converter (DAC) with a nominal output voltage range of ±10V and 16 bits resolution. While changing the reference voltage of the DAC allows to modify the output voltage range for all channels together, the DAC also has digital registers for setting an offset and gain value for each of its channels individually. This allows a high flexibility for selecting the range for the 16 *V_In_* (control voltages driving the current pumps) after production, and thus allows to tune the output current range of the current pumps to the actual experimental situation without sacrificing the 16 bit resolution. With the selection of this DAC, we obtain a natural partition of the scalable system into 16-channel blocks. We were able to obtain an update rate of 40k samples per second(25*μ*s time-resolution) for every individual channel.

For controlling the DAC, we use a Microsemi low-power nano FPGA (IGLOO AGLN250 VQ100) as delegate FPGAs. This very cheap FPGA showed itself capable of handling simple tasks with up to 25MHz. It does not need complex support components because it already contains flash memory for self-programming after power-on. It also comes with an IC package that can be used on two layer boards and easily be hand-soldered if necessary. In the supplemental materials section we included the design files for a board using the described FPGA and DAC, which was also used for our tests.

In figure 5, the structure of one 16-channel building block is shown. It is assumed that an external ‘master’ FPGA with connection to a PC provides data (with 25MHz over a 8 bit wide data bus) to such a building block about how and when to stimulate. In return, this master FPGA will receive feedback about error states (temperature warning, buffer over/underflows, etc) and thus be able to inform the user about putative problems.

**Figure 5:**
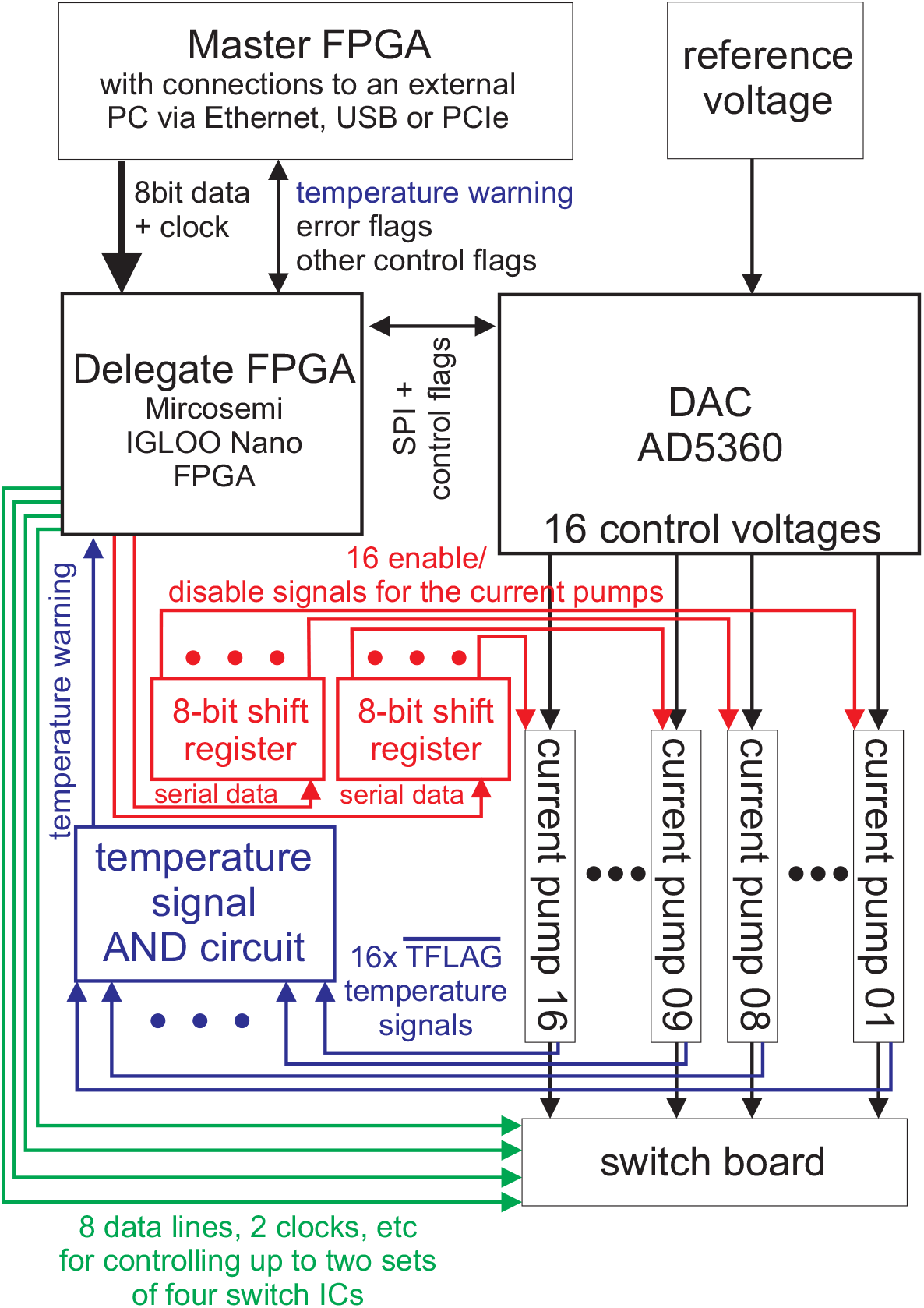
Sketch of a 16 channel stimulation building block. The building block receives data about the stimulation time series from an external ‘master’ FPGA (controlled by an external computer via Ethernet, USB, or PCIe) outside of the building blocks. The master FPGA provides the necessary information (clock signal, reset line, etc) and keeps all the blocks in synchronicity. One cheap ‘delegate’ FPGA for every 16 channel building block handles the incoming data and is responsible the timing of the stimulation. The delegate FPGAs program the digital-analog converters (DACs) via the serial peripheral interface (SPI) when the output of the DACs needs to be updated. Each DAC delivers the control voltages to its 16 current pumps which produce the stimulation currents. The delegate FPGAs also control if a current pump is enabled or disabled. For saving precious I/O pins of a delegate FPGA these signals are mediated by two high speed 8-bit shift registers with latched outputs. The temperature warning signals are reduced to one combined signals via AND gates (see figure 6) and then feed to its delegate FPGA. Putative problems are reported by the delegate FPGAs to the master FPGA. Furthermore, each delegate FPGA controls a variety of analog switches which protect the electrodes, the brain tissue, and additional measurement electronics.

The delegate FPGAs will check the data received from the master FPGA for inconsistencies because it is important to make sure that the intended stimulation signals are created correctly at the right time (in the firmware section, this topic will be discussed in more detail). Further responsibilities of the delegate FPGAs are to program the DACs via SPI interface, to enable/disable the current pumps when required, to control all other protection switches (if present, details in the following subsection), and to constantly check the temperature flags of the op-amps and if necessary, take preventive actions as a precaution for overheating.

In a 16 channel block, there are 16 signals for the current pump’s temperature warning flags and 16 signals for enabling/disabling the individual current pumps that need to be handled. Since each delegate FPGA has only 68 I/O pins and many of these pins are needed for other tasks, it is necessary to reduce the number of lines for checking the temperature and controlling the activity status of the current pumps. Figure 6 shows how a cascade of AND gates (e.g. SN74HC21PW) can be used to easily reduce the number of temperature warning lines from 16 to 1. With respect to enabling/disabling the current pumps (see figure 5), shift registers with latched outputs (e.g. two 8-bit shift registers SN74AHC595PW) can be used to reduce the number of connections from 16 to 4 (two data lines to the shift register ICs, and one line each for the shift register clock and storage register clock). It would even be possible to daisy-chain the two shift register ICs, thus saving one additional connection. Although it would double the time for programming the shift registers, this is not a problem if the AHC series of the shift register is used, because programming with 25MHz is easily possible. However, this additional reduction of control lines is not necessary for our design.

**Figure 6:**
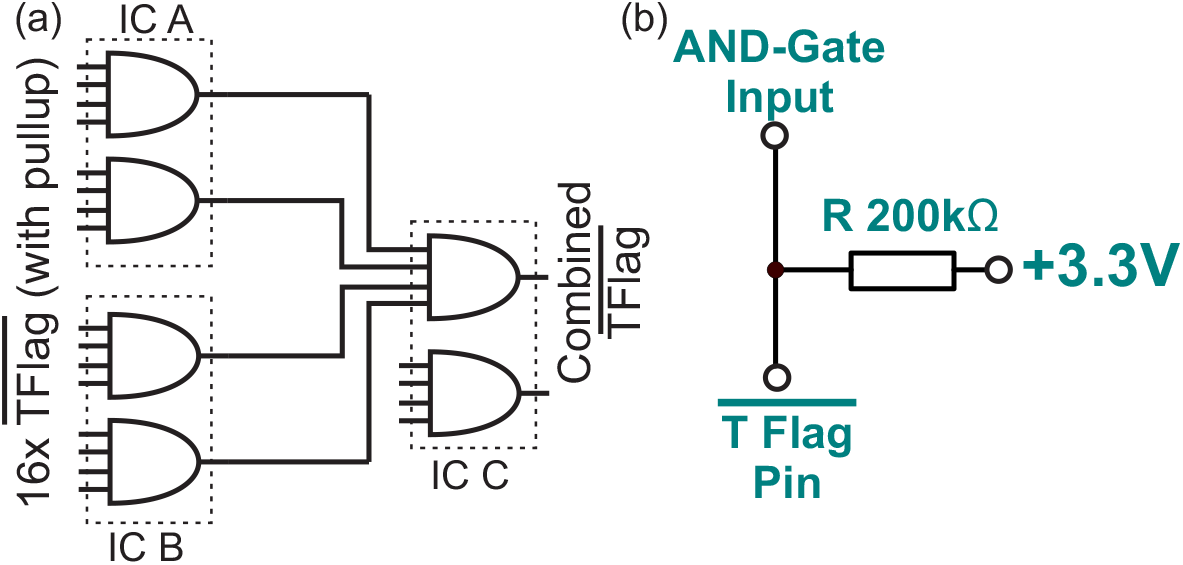
(a) Schema of how to reduce the required number of pins for the temperature warning flag using dual 4-Input AND Gate ICs (SN74HC21PW). (b) The temperature flag pin of the LTC6090 op-amp is an open drain output. For connecting it to an input of the AND gates, a pull-up resistor is required like it is described in the op-amp’s manual and is shown in this circuit diagram.

The schematics shown in figure 5 was transferred into an example design (design files are part of the supplement). The upper panel of Figure 7 shows the example design for a 16-channel current pump controller board. The four-layered board has a size of 89mm x 228mm and is loaded with components on its top side. The large area of the board is a direct consequence of the 0.75mm conductor spacing required for the high voltages (ΔV =140V). The current pump modules are plugged into the 16 arrays of female pin header connectors. This allows to easily replace broken modules as well as creates a radiator-type construction that allows a convective airflow for efficiently cooling the modules. As a side-effect, the use of dedicated modules makes it possible to use a four-layered board instead of a large six-layered board, thus saving costs.

**Figure 7:**
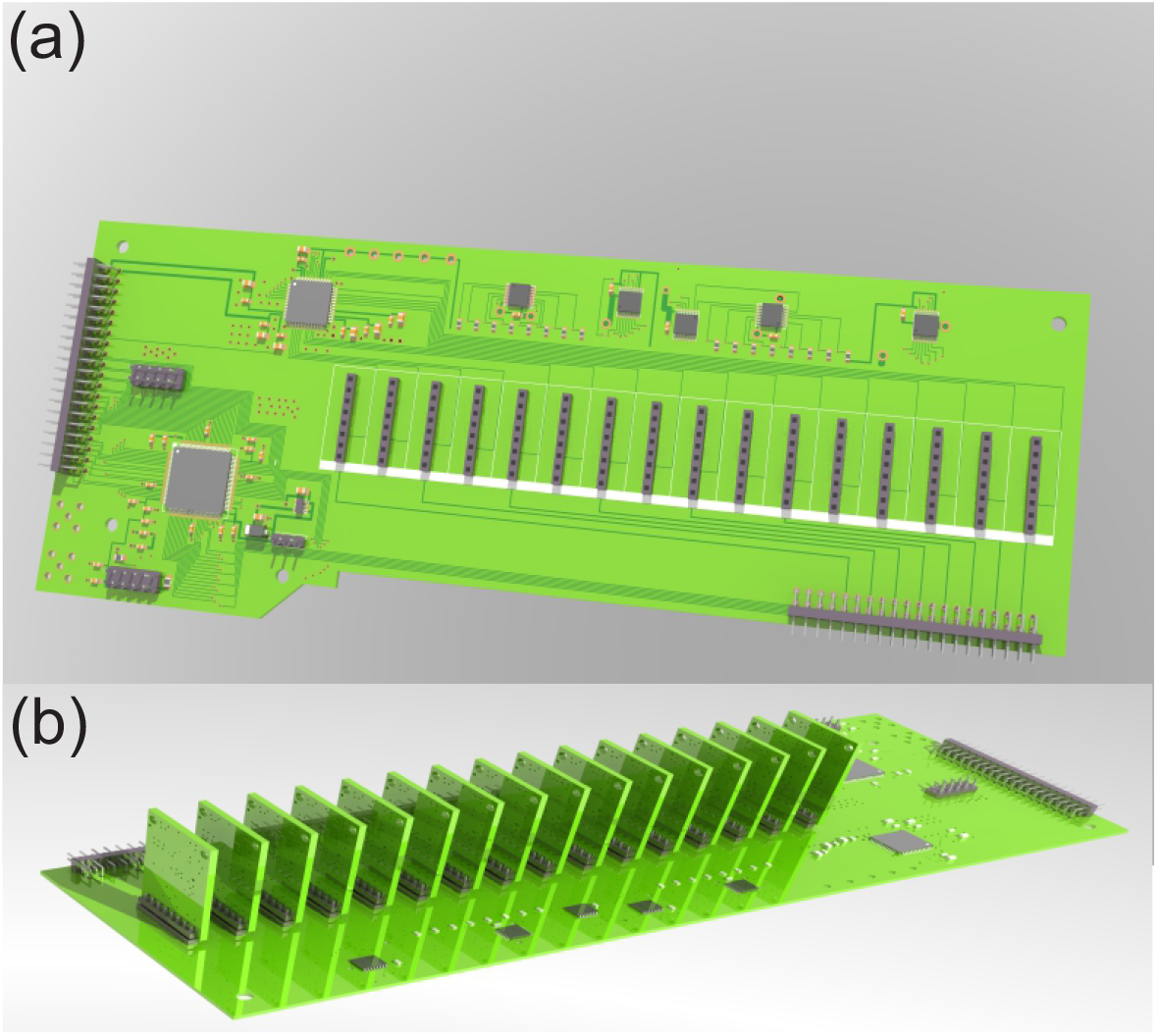
3D rendering of the example design for a 16 channel current pump controller board (89mm x 228mm, four layer board with single side component load) for Δ*V*=140V (The software couldn’t show the cooper planes for ground and the power rails which creates there the illusion of large unused areas). (a) The left connector carries the power supply, the reference voltage for the DAC, and the digital lines from the master FPGA (with the connections to the PC). In the lower left part of the board, the delegate FPGA is situated. The DAC is positioned in the upper left region. Right of the DAC, the shift register and AND gate ICs are placed. The lower right connector delivers the stimulation currents, digital control signals and power supply for analog switches (located on an additional switch board). A simple ribbon cable can be used for connection boards. As a result of the required electrical clearances due to the high voltages, the major area of the board is consumed by the 16 female pin header connectors. These are spaced far enough such that the metal part of components from one module can not touch the neighboring module as well as allow an airflow for cooling the current pumps. (b) Controller board with installed current pump modules.

We thought about adding digital isolators (e.g. Texas Instruments ISO764xFM) for separating the shift registers, AND gates and DACs from the parts of the circuit with high frequency digital signals. The purpose of adding digital isolators is to prevent distribution of potential high-frequency ‘noise’ over the system. Since we do not have the necessary equipment for quantifying the influence of our system on the neuronal signals we intend to measure, however, we postponed adding these isolators. Without the ability to measure, it is unclear if there really exists a problem, and if it can be fixed by adding isolators.

Another important issue is to choose a power supply with a clean ±70V DC power rail. This requires special solutions, which preferentially are suited for medical applications (even if they will first be used for animal experiments). We found one possible power source in the Vega750 system from TDK Lambda, where two +35V modules and two-35V modules are combined to create a ±70V supply. For improving the quality of the DC voltage, a Π-filter can be applied.

### 3.3 Analog protection switches

Simultaneously stimulating and measuring is a challenge. A measurement system for (extra-cellular) neuronal activities like the Intan Technology RHD2132 is designed to measure voltages in the sub-mV region. In the case of the Intan RHD2132, voltages outside of the range of ±400mV will be directed to ground via electrostatic discharge (ESD) diodes for protecting the analog inputs. For short time intervals with active stimulation current flow, the ESD diodes may protect the sensitive inputs for measurement from damage even if high voltages as e.g. 70V are used. For a longer stimulation duration this solution will not work, because the integrated ESD diodes will heat up rapidly and be destroyed quickly (in a worst case scenario, the diodes have to dissipate 25.6mA@70V=1.8 Watts). Another problem using ESD protection diodes is that they will consume part of the stimulation current if their threshold is reached, hence causing a lower current injected into the brain tissue than intended. Furthermore, overloading the op-amps in the measurement system with high voltages causes a large increase in their recovery time. During recovery, the op-amp will produce measurement artifacts lasting from a few milliseconds up to 100ms (or even more) while the timescale of interest for the neuronal signals lies roughly in the order of 1/10ms (action potentials) to 1ms (local field potentials). These two effects make it important to protect the analog inputs of the measurement system from stimulation signals with too large voltages. Another requirement for protection is that the current pumps must be disconnected from the electrodes when they are not used for stimulation (see subsection ‘Current Pumps’). If then the current pump is not disconnected from the electrodes, even if the DAC is programmed to a control voltage of 0V, small DC currents can be produced. This can be a result of mismatching resistor values or imperfection of the DAC calibrations. Furthermore, it blocks stimulation currents from one current pumps from flowing into others. Otherwise these currents are lost for stimulation.

The described problems make it necessary to include additional protection into the design. Figure 8 shows the solution we applied: During stimulation, the direct connection between the electrode and the measurement device can be separated by opening switch B. Current will then flow from the stimulator’s output through a closed switch C to the electrode (Fig.8(b)). After stimulation, switch B will be closed and switch C will be re-opened. Switch A, which was already closed during stimulation, now allows the electrode/brain tissue interface, which can act like a capacitor during stimulation and store charge, to discharge (Fig.8(c)). Finally switch A is opened and the system is again ready for measurement (Fig.8(d)).

**Figure 8:**
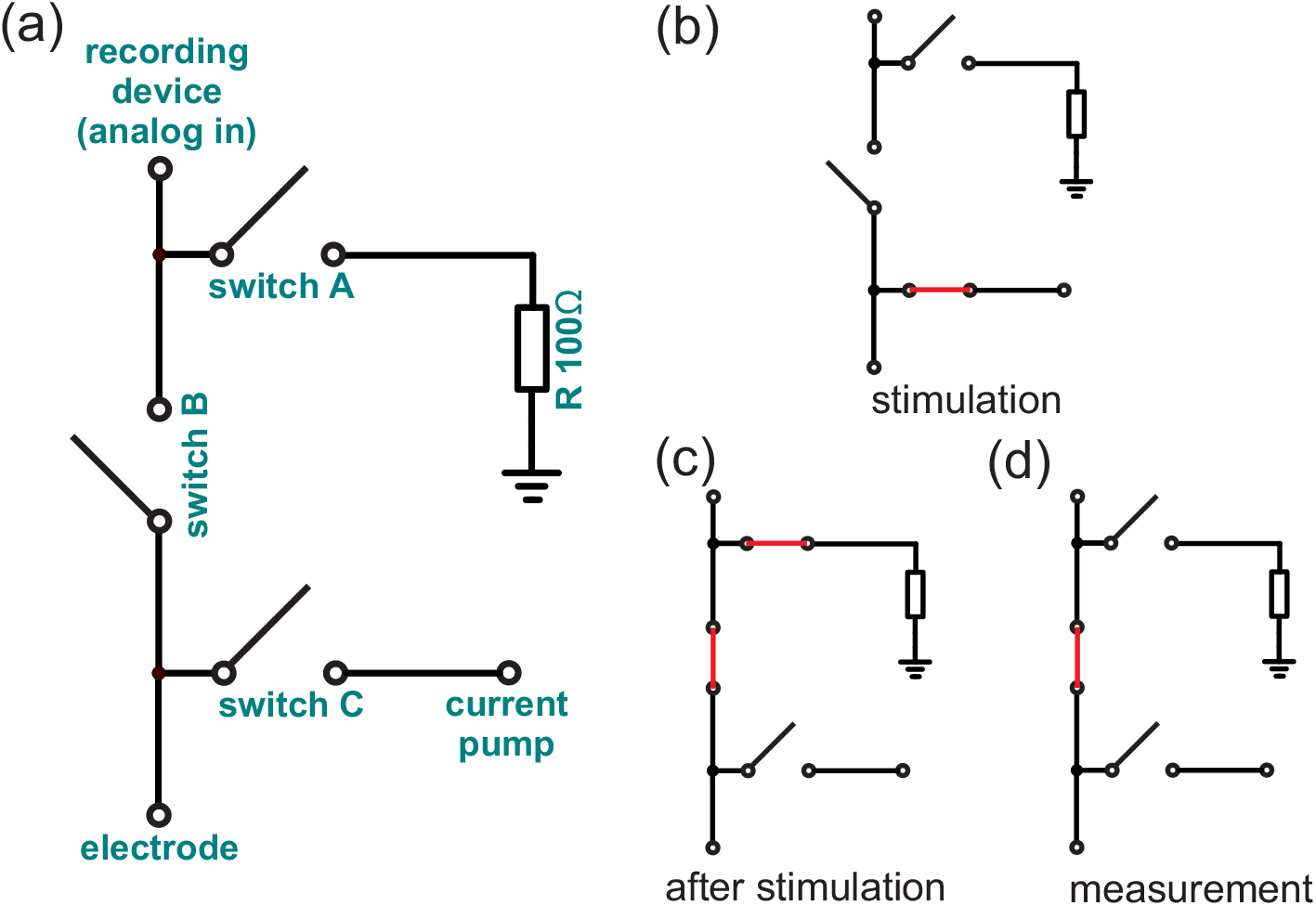
(a) Circuit diagram for protecting the analog inputs of a measurement system. (b) During stimulation switch B is open and switch C is closed, allowing the current to flow from the current pump to electrodes while no stimulation current can flow into the direction of the recording device. (c) After stimulation switch A and B are closed and switch C is opened. This removes the current pump from the electrodes and allows remaining charge (e.g. from the interface between electrode and tissue) to be bleed off over the bleeder resistor R. (d) Before resuming the measurement, switch A is opened.

In addition it is possible to add optional external protection diodes. These rather large discrete diodes, which can typically handle up to several Watts, can be used to unburden the ESD diodes of the measurement system. These diodes need to be fast like e.g. Schottky diodes and, obviously, need to be suitable for the properties of the neuronal signal passing by. If these diodes are installed, it is important to simulate (or test) the interaction between these extra diodes and the internal ESD diodes of the measurement system in advance.

For selecting the switches, it is important to consider the following constraints: Is the switch able to work reliably in the required voltage range of ±70V with the expected stimulation currents of up to 26*mA*? Which resistance does it add to the circuitry (should be small compared to the impedance of the used electrodes)? Does it have a signal transmission bandwidth of at least ≈50kHz? Is the time required for changing the state of the switch sufficiently small, e.g. 0.5ms or less? How often can the state of the switch be changed before it reaches the end of its lifetime? And very importantly, how much noise is added to the very small physiological signals passing the switch during recording?

Fulfilling all of these requirements, especially for voltages with an amplitude of ±70V, is a challenging problem for which we have not found an optimal solution yet. Electromechanical relays are too slow, requiring about 10ms per state change, and their lifetime expectancy is only ≈ 10^7^ state changes. Reed relays are faster (≈ 0.5ms) and their life expectancy is higher with up to 10^9^ switch cycles. However, assuming one state change every 10ms, 10^9^ state changes are reached after 2777.8 hours. If the system will be used eight hours every day, the relays will be worn out in less than a year. Solid state relays, a combination of a LED and a photosensitive MOSFET component, change their state with approximately 1ms and thus might be too slow. Pure MOSFET switches are also a possibility. Due to the positive and negative voltage load it is necessary to use at least two MOSFETs (and two diodes, if the internal diodes of the MOSFET are not suitable for handling the occurring stimulation currents). For operating these MOSFETs, a gate driver is needed that works in this high-voltage regime, thus adding to the list of components required for every individual channel.

While MOSFET-based switches seem to be a viable approach, we instead propose high voltage analog switches since they are easier to use. For our tests we decided to use the Microchip HV2201, which is an eight channel switch IC capable to switch analog signals with up to ±100V. These switches add ≈ 30Ω and have a turn off/turn on time in the order of 5*μ*s. In our test environment the switches worked successfully, but we found that two of the eight channels broke by handling them during our tests. As a consequence, we modularized the switch ICs in our example design (see figure 9(a)), allowing easy replacement in case they fail. Thus the switch modules only host the switch ICs, while the switch board realizes the circuit diagram shown in figure 8. Although a 16-channel version of this IC is also available (Microchip HV2601), we recommend to use the eight channel version because it is easier to obey the required 0.75mm conductor spacing for Δ*V*=140V due to additional free space by internally non-connected pins and the cost per channel is the same.

A remaining question is which noise influence the switch IC has on the recorded signal. Sadly, we were not able to answer this question because our lab does not have the necessary specialized equipment to generate and measure such small signals reliably. In cases where high stimulation voltages are not needed, and hence the gap between the maximum voltages of the stimulation and recorded signal is smaller, there might be an analog switch IC solution which might have a lower noise influence. Also reducing the supply voltage to its minimum may improve signal-to-noise ratio. In the worst case, where the noise behavior is not acceptable, only the switch modules need to be replaced by a low-voltage solution while the larger four-layer switch board (see figure 9(c) and (d)) can still be used.

**Figure 9:**
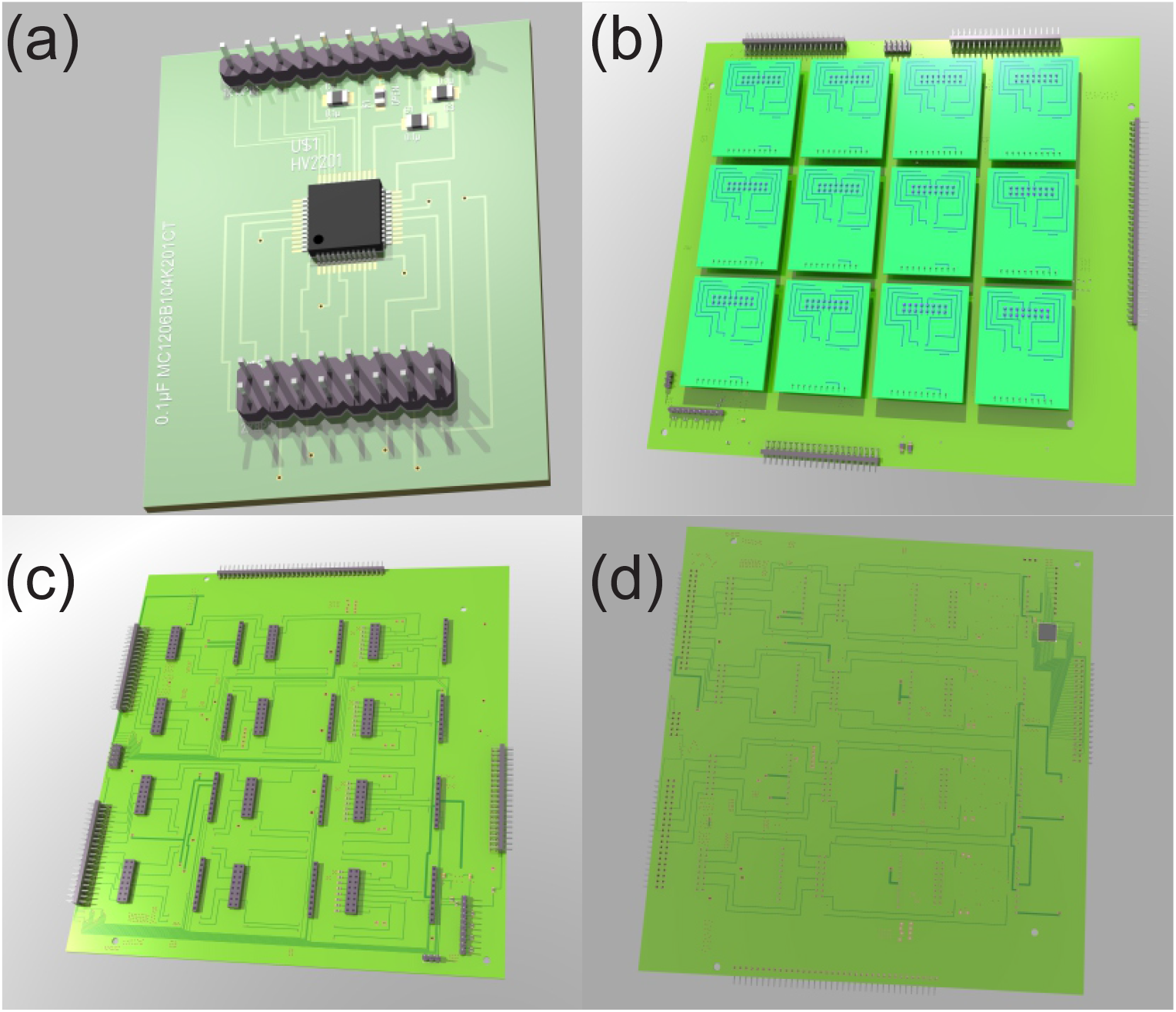
3D rendering of the boards used for switching the analog signals and currents. The boards are designed for a for Δ*V*=140V (The rendering software couldn’t show the cooper planes for ground and the power rails. In this pictures this creates the illusion of large unused areas.). (a) Module hosting the switch IC. This 38mm x 49mm two layer board with single side component load, contains a Microchip HV2201 8 channel high voltage analog switch. The pin header with 1 row of pins connects to the digital signals and the supply voltages while the other pin header is concerned with the analog signals. (c) Four layer board (209mm x 197mm) realizing the electrical circuit shown in figure 8 for 32 individual channels. (d) While most of the components are on the top side, an optional 32 channel Intan RHD2132 can be installed on the flip side of the board. This would provide the necessary parts for measurements, in an OpenEphys-compatible fashion, directly on this board. The two pin header on the left are used to connect two of the 16 channel current source board (see figure 7) via ribbon cables. These connectors deliver the required power supply, digital control signals for the switches, and the stimulation currents. It is important that these ribbon cables have a low capacitance. Otherwise (e.g. in the case of coaxial cables) the biphasic stimulation pulse might be absorbed by the cable’s capacitance. On the upper position on the board, the connector for electrodes is situated, while on the right side a connector for a recording system is available (if the Intan RHD2132 is not used). (b) Switch board with installed switch modules.

### 3.4 FPGA firmware

Besides the hardware, the system needs to organize the distribution of information and manage the operation of its components. We decided to use FPGAs for ensuring a precise timing of the stimulation sequences. One external, master FPGA establishes the connection to and from a host computer (e.g. over a fast Ethernet interface) and communicates with multiple ‘stimulation modules’ over a shared bus. Each stimulation module comprises a 16-channel stimulation building block which includes a delegate FPGA (Microsemi IGLOO nano). The delegate FPGA’s job is to program the DAC, the 8-bit shift registers and the other input protection switches as well as monitor the overheating signal from the current pumps. In the following, we will describe the firmware of the FPGAs and the performance of the system in detail.

#### General design aspects

For realizing a particular stimulation time series, two approaches are possible:

a. Stimulation patterns are defined before the experiment is started, and stored in a memory bank of the master FPGA. During the experiment, the host PC can then instruct the master FPGA to replay the sequence from its memory with precise timing.
b. The firmware is designed in the spirit of a ‘continuous data stream’ process. The stimulation data is fed continuously into the system and the delegate FPGAs process this stream of instructions and act on it as soon as possible.

In our test system, we realized option (b) since it is more flexible. First, it allows arbitrary stimulation patterns to be generated by the host PC, sent to the master FPGA, and distributed to the delegate FPGA’s for execution. Second, it allows to extend the stimulator to a closed-loop system capable to adapt the shape of the stimulation pattern to the measured neuronal activity state in real-time.

#### Data transfer between master FPGA and delegate FPGA’s

For keeping the amount of physical connections between the stimulation modules and the master FPGA (with the connection to the PC) low, we decided to implement a broadcast approach for the communication. Multiple stimulation modules listen at the same physical data bus originating from the master FPGA. The master FPGA sends out all the stimulation information with a clock of 25MHz using a bus width of 8 bit, thus equaling 25 MByte/s per bus.

From this common broadcast stream, the individual controller boards filter out the messages concerning them. Every individual nano FPGA firmware is compiled with a 7-bit identifier, which allows up to 128 controller boards to be addressed. Including a header for determining the beginning of the data package and two bytes reserved for future extensions (e.g. check-sum for protection against data corruption), 56 bytes are required to describe the new state of a 16-channel block (table 2), namely:

**Table 2;.** Structure of the data package

|  |  |
| --- | --- |
| 2 bytes | header |
| 1 byte | address of controller board<br>(7 bits=128 boards, 1 bit for broadcasts to all boards) |
| 36 bytes | new state of DAC |
| 10 bytes | new state for protection switches |
| 5 bytes | duration of new state |
| 2 bytes | reserved for extensions |
| 56 bytes | in total |

For every state change, the current pump controller board requires 18 bits for every channel of the DAC, consisting of a 16-bit data field plus 2 bits for mode selection. Typically, the data field holds the control voltage, but can in conjunction with the mode selection bits be used for trimming offsets and gains of the individual DAC’s channels. In total, 18 bits times 16 channels sum up to 36 bytes for defining the new state of the whole DAC. Each state change also requires 10 bytes (5 bits times 16 channels) for defining the state of every channel’s four individual protection switches (4 bits) and the corresponding on/off control (1 bit) for their current pumps. To reduce the amount of data transferred for realizing a stimulation time series containing constant segments or for pausing stimulation, every message packet contains one 40 bit value specifying the duration of a new state which can last up to 4500s. The delegate FPGA will hold the new state (i.e. switch and current pump settings, stimulation currents) for the required duration before the next set of instructions will be processed. Every message received during that time will be buffered in a FIFO.

Aiming at a maximal update rate of 40kHz, one 16-channel block requires a data rate of 2.24 MByte per second. This allows one bus to easily control a system with up to 176 stimulation electrodes (11 x 16-channel blocks). If higher update rates and/or larger number of stimulation electrodes are required, the external FPGA could easily manage and synchronize several broadcast buses.

#### The connection to the outside world – the Master FPGA

The firmware of the master FPGA (a Xilinx Spartan 3A on an Orange Tree Tech ZestET1 board with 1GBit Ethernet connection, www.orangetreetech.com) can be kept very simple. In our test implementation it takes data from the PC, buffers it into a FIFO, and then broadcasts it to the delegate FPGA’S on the controller boards. In the direction to the PC, only information about temperature warning status, error states and buffer status is transmitted back to the user. The corresponding test software for the PC is also very simple: It just builds data packages and switch states based on the desired stimulation time series and then transmits it via network stack of the operating system to the master FPGA. Both FPGA firmware sources and the PC test software are part of the supplement.

#### Latencies and Synchronization

If the stimulation system is controlled by a PC, which is not a real-time device and thus occupied with other tasks in parallel, some considerations have to be made how to synchronize stimulation with external events. These could be the onset of a visual stimulus shown on a screen, or the start of an external recording system. As long as the FIFOs of the delegate FPGAs contain a sequence of stimulation instructions, execution is deterministic and in synchronization with the clock signal provided by the master FPGA. Hence for longer stimulation sequences, the task of the controlling PC is to segment the sequence into appropriate instructions, transmit them to the master FPGA, and keep the FIFOs well filled for bridging time intervals where the PC is occupied with other duties (e.g. interrupt handling). Obviously, there is a balance between the necessity of keeping FIFO buffers filled and the possibility to flexibly adapt stimulation to external events.

For synchronizing the FPGA clock with an external device, the master FPGA can be disconnected from the internal clock source and connected to an external clock input. In addition, we provide a ‘pause’ input which holds execution of valid messages in the FIFOs of all delegate FPGAs when the corresponding pin is set to low (digital ground), allowing to execute a stimulation sequence in synchrony with an external event: First, the controlling PC first has to make sure that no stimulation commands are pending. This can be done by querying the master FPGA. Then, the external ‘pause’ input must be set to low holding execution of stimulation messages, after which the intended stimulation sequence can be programmed by the PC. When this task is finished and the external event happens, the pause input is set to high (+3.3V) thus resuming delegate FPGA operation and immediate execution of the programmed stimulation sequence. Note that this functionality only provides a basic mechanisms for synchronization – for realizing more sophisticated schemes, the firmware of the master FPGA has to be extended accordingly and to the needs of the intended application.

For an application requiring a less precise timing, it might be sufficient to ensure that the FIFOs are empty, and then to transfer the stimulation sequence from the PC directly. For modern PCs with Gigabit Ethernet connections, we expect latencies below 1ms (the actual roundtrip time can be easily accessed within the test implementation).

### 3.5 Tests

Besides the design of the system concept and the layout of all boards for an example production system, we designed and built test boards (PCB design files can be found in the supplement). Figure 11(a) shows this setup. Its goal was to test if the single components worked as intended, test their interplay, and debug the FPGA firmwares and PC software as well. As a result we obtained a fully functional system (tested with a supply voltage of ±15V), laid out for 16 channels, but populated with current pumps and switches for only two channels. Stimulation sequences could be sent successfully by the PC over the network to the Orange Tree Tech ZestET1 (Figure 11(b) and (c)). The ZestET1’s FPGA acted as master FPGA and distributed the information to the delegate FPGA which in turn controlled the current pumps and protection switches. At the output of the switches we placed a variable load, realized by a potentiometer, in series with a shunt resistor. We measured the voltage drop over the shunt resistor with an oscilloscope and found that the system and its current pumps showed the expected behavior under different loads (Figure 12). In-depth performance and noise measurements have not been done because the long conductor leads used in the setup picked up a fair amount of noise from different external sources.

**Figure 10:**
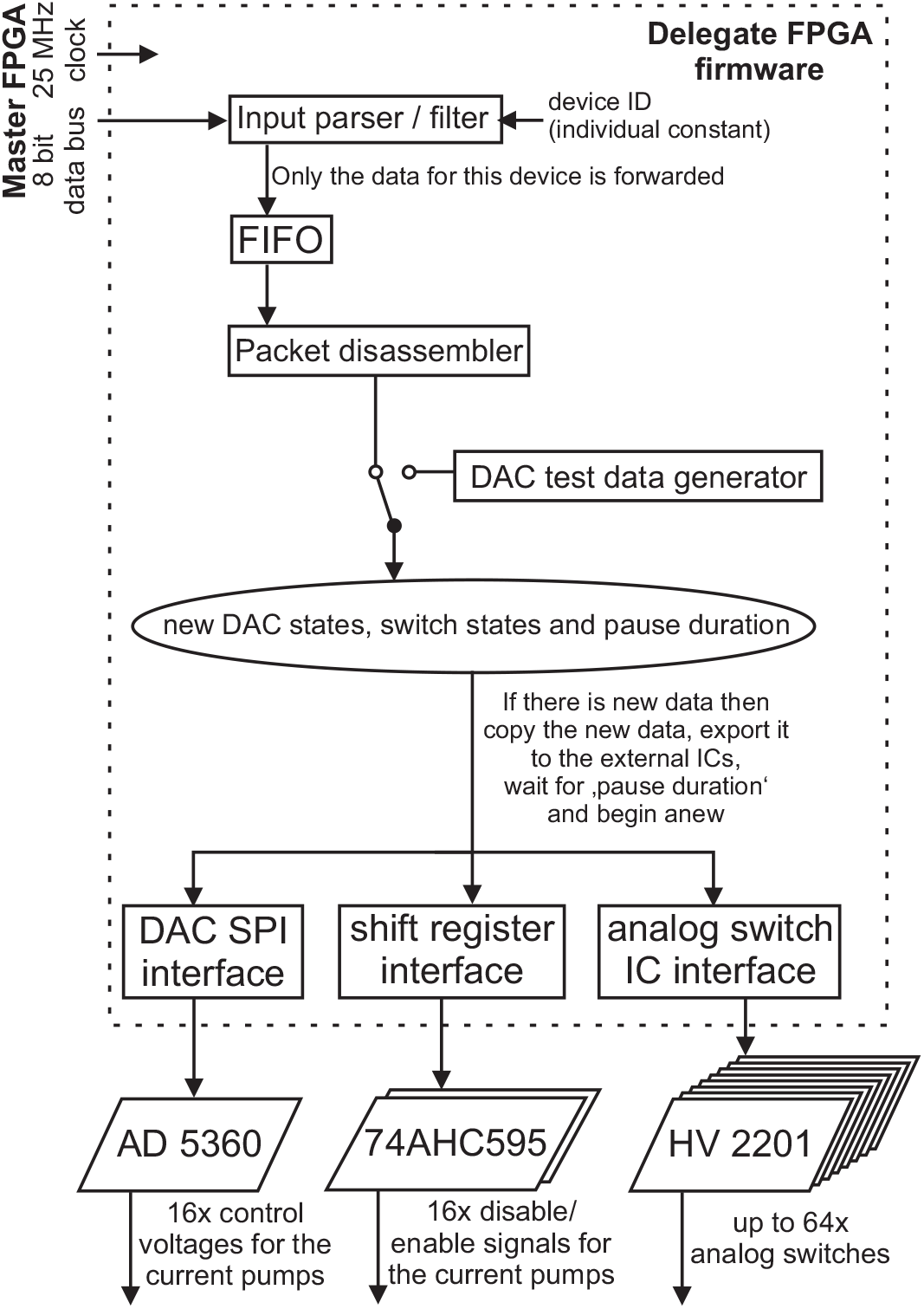
Simplified schematic of the firmware for the delegate FPGA. Data from the master FPGA is received over a 8-bit wide and 25MHz fast data bus. After power-up one 12 byte long entrainment block, is sent which makes sure that the data transfer between the FPGAs is working correctly. Then the nano FPGA receives 56 byte long data packages containing the stimulation descriptions. If the data package was signed with the address id for the delegate FPGA (which is individually programmed into each delegate FPGA firmware), the data is copied into a FIFO for further processing. If the address id is unknown then the data package is ignored. The accepted data is taken from the FIFO and is disassembled into a set of new DAC values, switch/ current pump states and a pause duration. When the rest of the system is ready for processing a new set of instructions, the disassembled instruction set is copied for further processing. After that, the package disassembler can continue and prepare a new set of instruction in the background. The copied set of instructions is analyses by three finite state machines (FSM) and transferred to the DAC (via SPI), the two 8-bit shift registers and up to eight analog switch ICs. After that, this part of the system waits, for the time defined by the pause duration, and then continues with the next set of instructions from the packet disassembler. Beside producing the control voltages for the current pumps, the DAC has an additional digital I/O pin (GPIO) which is connected to an input of the delegate FPGA. The logic state of this GPIO is flipped through SPI at the beginning of programming a new set of instructions into the DAC. This allows to observe the timing of the DAC. Also a test data generator is included in the firmware, which can generate a continuous saw tooth voltage signal at the DAC’s output channels. This signal generator can be enabled during compiling the firmware.

**Figure 11:**
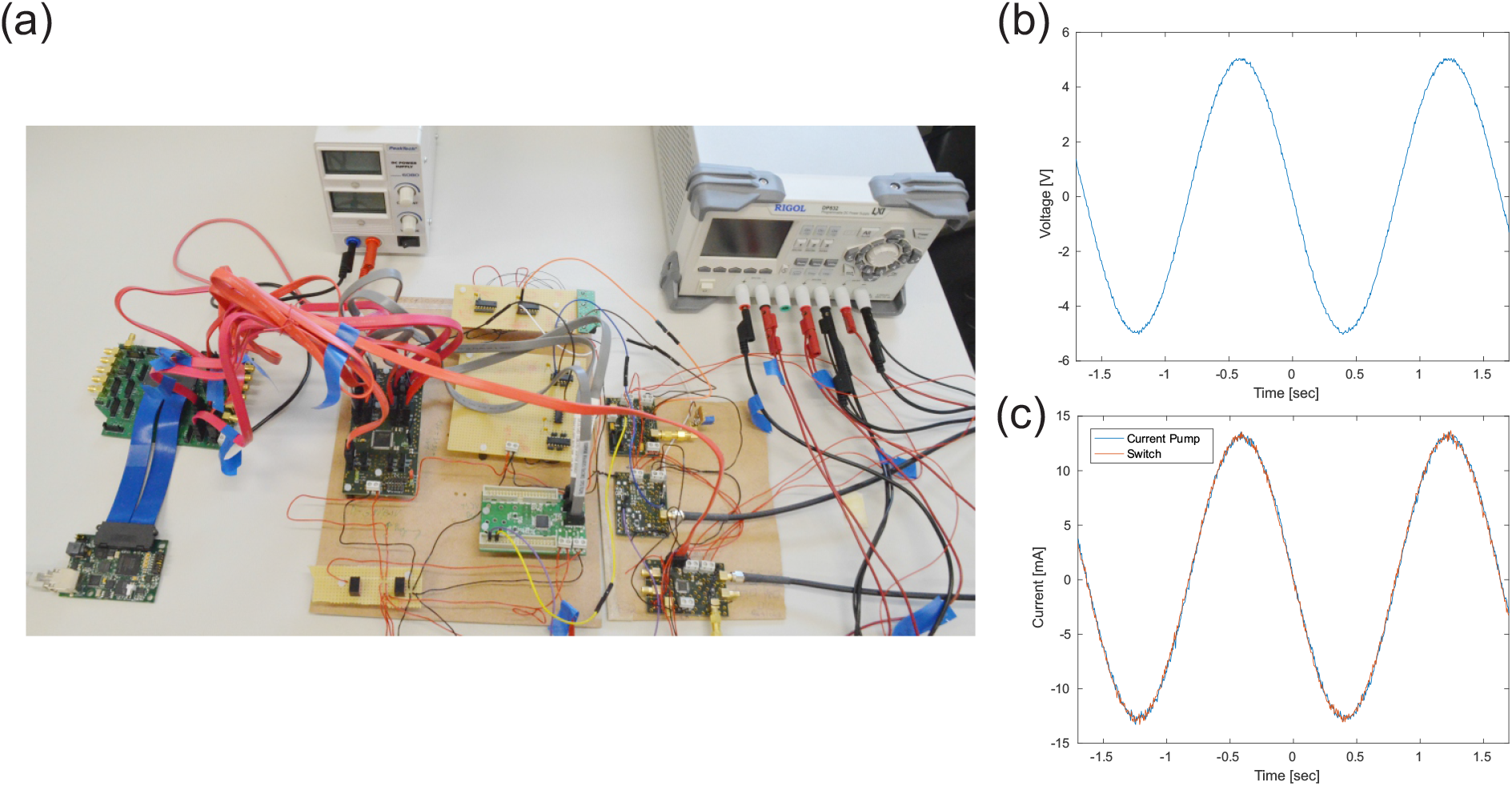
(a) Used test setup for testing the system concept and debugging the firmware. The setup showed that the concept behind the firmware, software and hardware works as intended. (b) Voltage output of the DAC created by a timeseries of 65535 points for one oscillation. (c) Corresponding current output to (b) measured after the current pump as well as the analog switch. The resistive load was 52 Ω.

**Figure 12:**
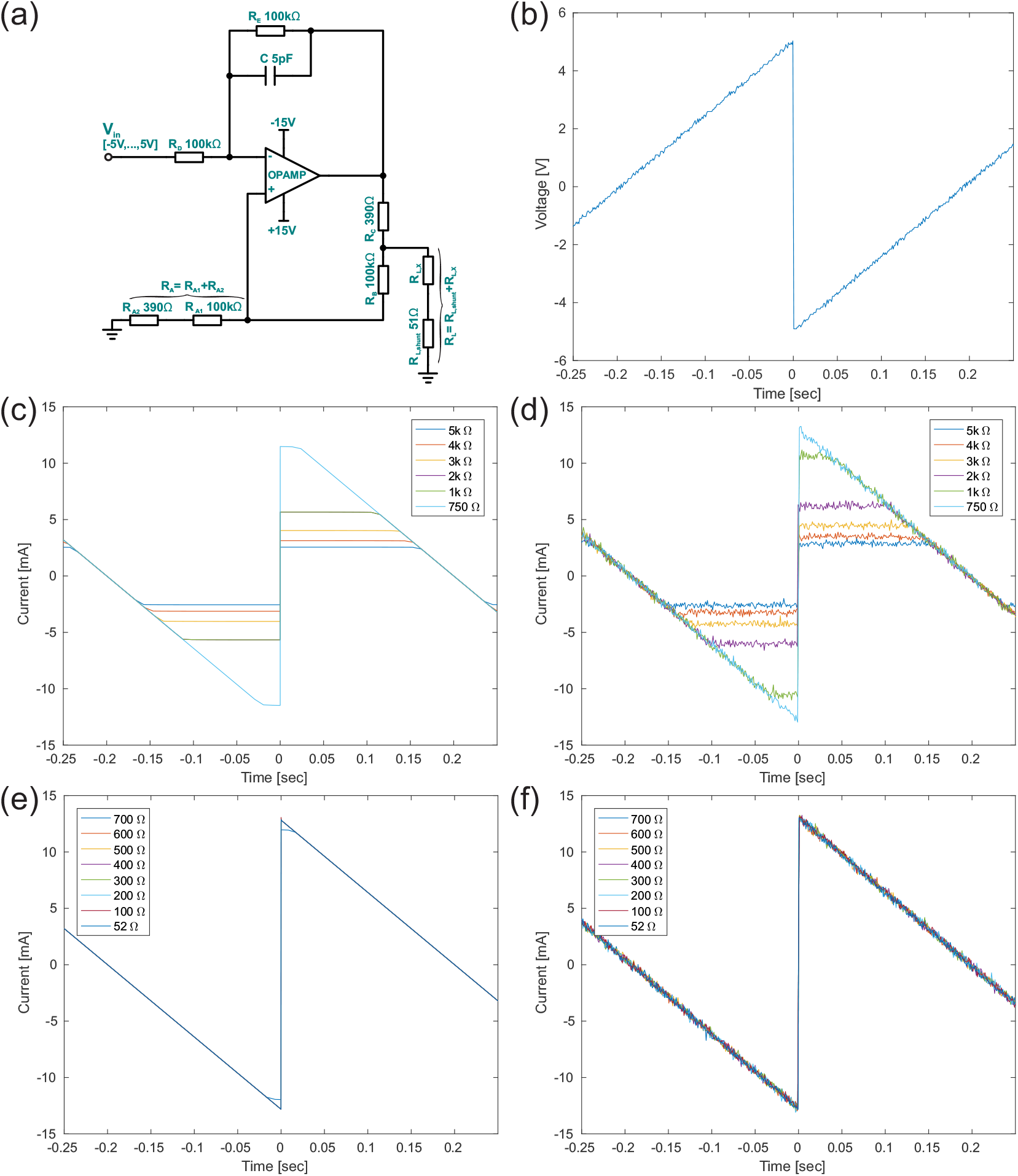
(a) Circuit diagram of the test setup. The delegate FPGA’s test pattern generator
was used for producing a test signal. It rans continuously through all the possible 16-bit integer values of the DAC. The output current of the current pump was measured via the voltage drop over the shunt resistor. The resistive load *R_L,X_* was varied. The supply voltages were ±15V. (b) Voltage output of the DAC. Due to historical reasons, the reference voltage for the DAC was only 2.5V which is half of the recommended reference voltage. As a result the output range of the DAC is reduced to a range between −5V and 5V (instead of ±10V). Furthermore, the halved output range of the DAC and a supply voltage of only ±15V results in the current pump to produces only half of its maximum current output. (c) Simulation of outputs for *R_L_s* that can’t be driven by the given supply voltage. (d) The corresponding measurements to (c), recorded with a digital oscilloscope (Agilent InfiniiVision DSO 6102A). (e) Simulations similar to (c) but with smaller *R_L_s*. (f) shows the measurements for the curves shown in (e).

## 4 Discussion

We presented a system concept for a multi-channel stimulator with protection to be used in parallel with a measurement system. The main design goal was to enable the measurement system to restart recording as quickly as possible after a stimulation pulse while protecting its delicate analog inputs from the high voltages produced during stimulation. This feature is especially important for real-time closed-loop applications such as cortical prostheses, in which stimulation has to be adapted to on-going brain activity and delivered with millisecond precision. The system is based on blocks of Howland constant current pumps with 16 independent channels controlled by one ‘delegate’ FPGA. These blocks can be easily combined to stimulate with several hundreds of electrodes simultaneously. The findings from developing the system concept was condensed into circuit board designs for driving ultra small surface electrodes which may require stimulation currents of up to 25mA per electrode generated by voltages of up to ±70V. Test boards with the components planned for being used in this system have been designed and built. Furthermore, firmwares for the FPGAs and software APIs for a controlling PC have been written. The feasibility of the concept was shown in simple test measurements.

Using 176 electrodes from 11 stimulation blocks attached to a single message bus in parallel, arbitrary time series of stimulation currents with 40k samples/s can be generated, covering the typically used shapes [78]. All design files are part of the supplement.

### Interfacing the stimulator: towards a closed-loop system

For controlling the stimulation blocks with the delegate FPGAs, two approaches are feasible. (a) a small FPGA just distributes the data it receives from the controlling PC. (b) one can use a more flexible solution with a large FPGA which can be used to take over tasks from the PC. The later solution provides the opportunity of realizing a closed-loop system.

Before using the Orange Tree Tech ZestET1 board, we tried to use the USB module FTDI UM232H with a FTDI FT232H IC first. The UM232H is one order of magnitude cheaper (≈ 30 Euros, the test board is part of the supplement). The idea was to connect the UM232H directly to the delegate FPGA. Hence each 16-channel block would have been controlled by its ownUSB link to the PC. Industrial USB switches based on PCIe cards with 48 USB ports or more are available and would ensure that this solution would scale to the number of several hundreds of channels. However, first tests indicate that this cheap solution has its own caveats: in the fast ‘FT245 synchronous FIFO interface’ mode the Microsemi Igloo nano FPGA (as the delegate FPGA) was not able to handle the necessary speed and timing for handling the enforced data-flow from the UM232H. We also tried the ‘FT245 style asynchronous FIFO interface’ mode, where the delegate FPGA controls the speed of the connection. Here, the timing worked out, but we measured one transmission error approximately every MByte of data transferred.

Instead of spending the time for debugging, we therefore switched to the more powerful ZestET1 for which we had in-depth experience from another project. This solution worked perfectly. Apart from the limited number of I/O pins the ZestET1 provides for buses to the delegate FPGAs, an important bottleneck for the broadcast approach lies in the data bandwidth of the connection between the ZestET1 to the external PC, which is responsible for the calculation of the stimulation time series. A solution is to perform the analysis of the recorded data and calculation of the pulse sequences directly on the ZestET1 (or a board with an even larger FPGA). This allows more freedom in organizing the flow of data to the 16 channel current pump controller boards and reduces the latency between measuring the actual state of a brain network and the intervention through stimulation. The external PC would then only be used to configure the stimulation system, and for non time-critical tasks. Besides reducing control effort and external data traffic, this extension also allows to realize a real-time closed-loop system suitable for medical applications.

### Optimizing switching logics

When operating a multi-channel closed loop system, we only want to disable recording for electrodes which are used for stimulation or disturbed by the stimulation pulses from neighboring sites, and keep the other ones recording. However, in a non-homogeneous medium like the brain/electrode interface, it is very problematic to estimate the extent and strength of the current distribution from one electrode spreading to the other electrodes during stimulation. One way to approach this issue is to add an additional ADC suitable for larger voltages, and to use it in a test phase for measuring the average current spread from stimulation at single sites, and from stimulation with typical sequences intended for regular use. These measurements would allow to determine which electrodes can be kept recording while stimulation is performed. However, the proposed extension would require an additional analog signal multiplexer (capable of handling such high voltages) for selectively connecting single electrodes to this ADC. Again this would rise questions about how much additional noise is introduced into the system by the added components, and how much stimulation current the multiplexer would consume itself.

### Optimizing and miniaturizing the system

Is it really necessary that a stimulation system has to provide voltages up to ±70V for delivering the currents needed to successfully communicate with cortical networks? Unfortunately, this question can not be answered yet. Resistance/impedance measurements of the electrode grids that we intended to use in our project [42, 77] show strong hints in this direction. However, these tests were performed with the electrodes in a saline solution, and they were conceived for understanding the electrode decay over time as well as for determining the best type of surface coating to be used. Measuring the real impedance of the electrode/ brain interface, and quantifying how it changes over time is a problem because such a measurement requires current flow between electrodes in a living animal. Furthermore, it is unclear how large the stimulation currents need to be for creating phosphenes (sensations of light blobs) by surface stimulation, and whether it is possible to produce stable phosphenes at all [29],[28]. Choosing too large stimulation currents will result in seizures [28], [47] or in the destruction of tissue and electrodes. These are important questions that need to be answered first, before this technology can be transferred into medical applications. For this reason we designed a versatile stimulation equipment which can be flexibly used with a variety of electrode configurations in different experimental paradigms, thus allowing carefully planned animal experiments for determining the optimal approach for an intracortical, bi-directional brain-computer interface. For a production system, this knowledge can then be used to narrow down the specifications for the stimulator to the relevant parameter range.

In the case that high voltages are not needed, the circuit diagrams do not need to change. Only the gain resistor values in the Howland current pumps have to be adapted. However, a much denser packaging of the components will then be possible. Figure 13 shows an example concept of a more compact construction while keeping its maintainability (broken components can still be fixed by simply replacing modules) as well as allowing for sufficient airflow through the construction for cooling.

**Figure 13:**
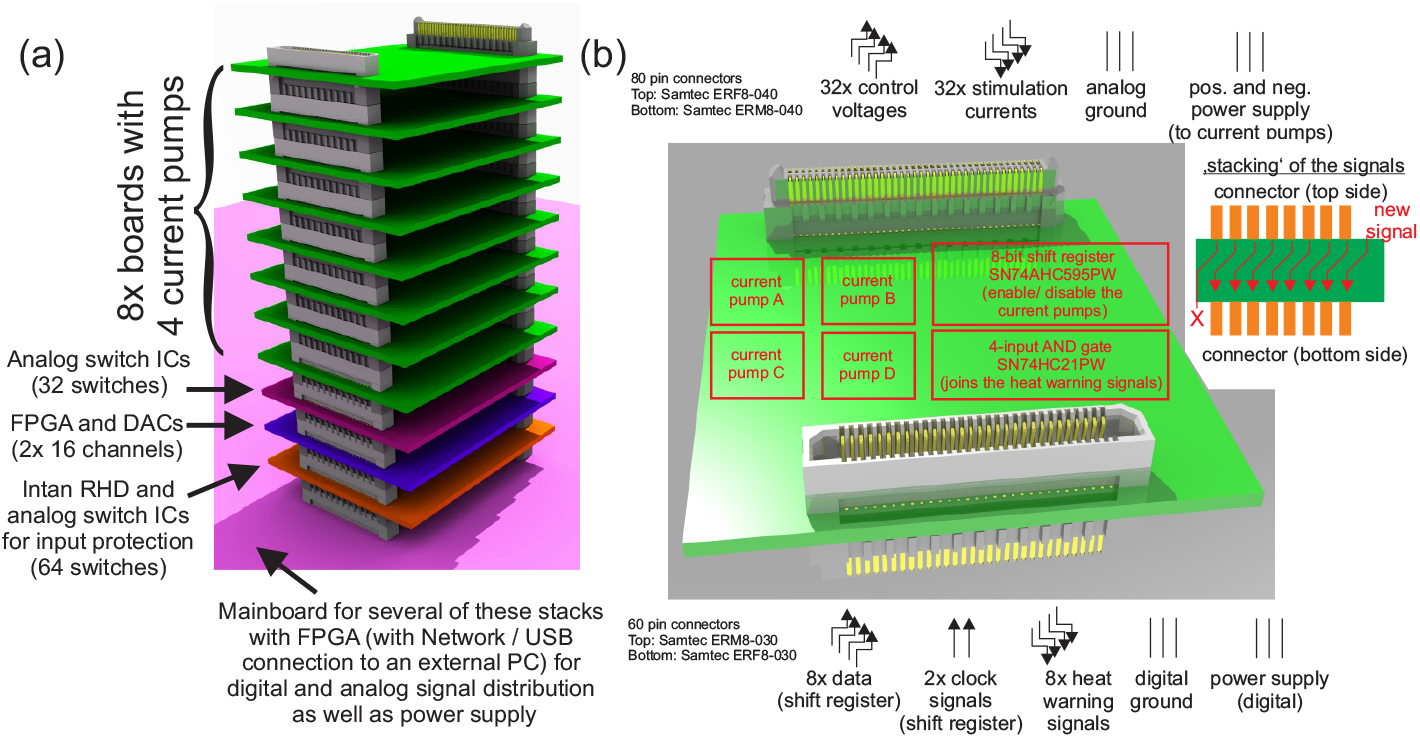
(a) Idea for a more dense construction for the stimulator if lower stimulation voltages can be used (or conformal coating can be applied). On a mainboard (pink surface), which is responsible for distributing the power supply and information containing the stimulation and switch states, stacks of several pagoda (only one is shown) are hosted. One of these pagoda are made of four different types of modules. On the lowest level, a module containing a 32 channel Intan RHD2132 (which could be connected to an OpenEphys recording system) with external analog input protection (two switches for each channel) is placed. On the second layer a nano FPGA and two 16 channel DACs are situated. On level three, analog switches can be found for disconnecting the current pumps from the lower rest of the circuit. From layer four up to layer eleven, identical modules with four current pumps each are installed. The pagoda has an approximately high of 12cm and uses the same combined circuit diagram as our example designs. (b) These modules, used from layer four up to layer eleven, consist out of four current pumps as well as a 8-bit shift register for turning the current pump on/ off and a 4 bit AND gate for joining the temperature warning flags of the current pumps. ‘Stacking’ of the signals (which is inspired by a bit-shift operation; see inset) allows to daisy chain the same identical modules over 8 layers for a total of 32 current pump channels.

Another opportunity for simplification would be to build a new design around the recently released Intan RHS2116, if the intended neuroscientific application fits into the specifications of that chip. This one-chip-solution is a 16-channel bio-signal amplifier with 16 bit ADC and programmable current sources/sinks based on a DAC with only 256 steps. The DAC can deliver up to 2.55mA with up to ±14V including some input protection. These specifications might fit if stimulation percepts such as e.g. visual phosphenes are to be created by intra-cortical stimulation instead of using surface stimulation [47]; an approach we believe to be better suited for long-term stable clinical applications. However, the input to the internal amplifier of the RHS2116 can not be disconnected from the current pumps or the electrodes, and it has a capacitor directly connected to the inputs of the op-amps which is charged during stimulation. This is not an optimal solution if the time between stimulation and continuation of the recording should to be as small as possible.

Further miniaturization may require to leave the realm of off-the-shelf components. Figure 14 shows an idea for an active adapter for Intan RHD2132 bare dies. This micro-machined Si adapter contains buried analog switches (e.g. [81]) to protect the analog inputs and allows bleeding off remaining charges over a bleeding resistor. These switches are controlled by shift registers with state memory which will also be embedded within the Siadapter. The Intan RHD2132 is wire-bonded (or flip-chip bonded) on top of the adapter. The flip side of the adapter is constituted as a ball grid array, allowing a much simpler assembly process compared to the normal bare die Intan RHD2132. A miniaturization of the current pumps will also lead to significant smaller stimulators [33].

**Figure 14:**
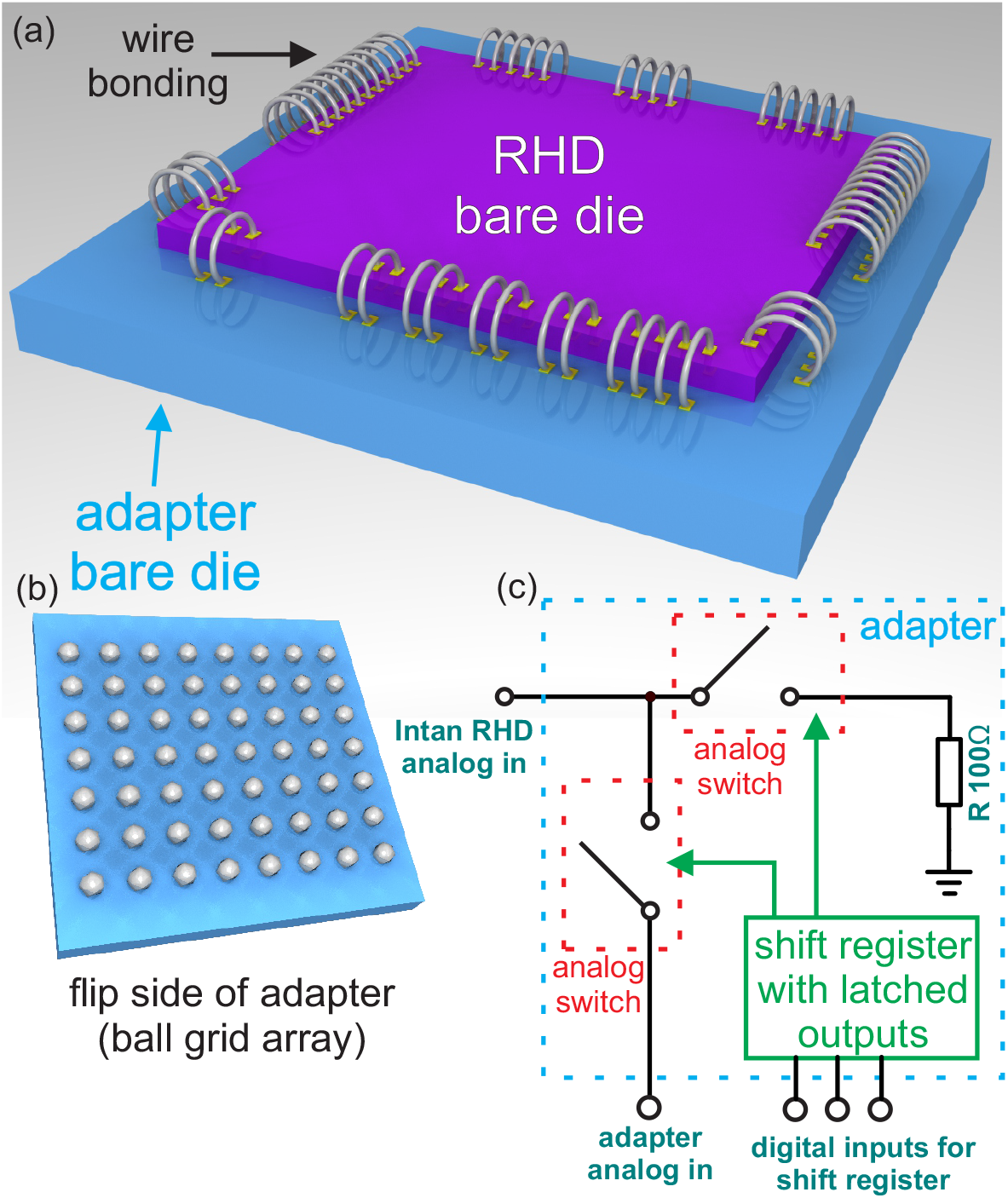
Idea for an adapter for fostering miniaturization. (a) This micro-machined Si adapter carries on its top a 32 channel Intan RHD2132 bare die for recording neuronal signals which is flip-chip or wire-bonded to the adapter. Inside of the adapter, every analog input to the RHD2132 is protected by two analog switches and a bleeder resistor ((c) circuit diagram of the input protection). Also a shift register with state memory (functionality like the SN74AHC595PW) is buried in the adapter for controlling these 64 analog switches with only a few pads. (b) The flip side of the adapter, which is used to connect to a circuit board, is shaped like a ball grid array allowing an easy assembly.

The ultimate goal in the future is to develop a medial implant allowing intracortical stimulation. The challenges of building such a fully implantable wireless system [82, 83, 84, 85, 86, 87], [88, 89] arise from the necessity to obey multiple constraints: the height is restricted, especially if the implant should be placed between the skull and the brain [90] for improving long term stability. Here, possible pressure to the brain tissue is an additional problem [91, 92, 93]. The maximum area available for the implant is limited by the curvature of the skull/brain and the rigidity of the implant. If the active parts of the implant are close to the brain, then heat produced from the consumed power is directly transferred to the tissue, bone and fluids [94, 95]. Therefore, it is necessary to keep the energy consumption low or otherwise it could result in damage to the body [96]. Further important problems are bio-compatibility [97] and long-term stability [98] since the implant needs to survive inside the body for many decades, and must not harm the brain tissue during that time [99, 100, 101]. Furthermore, for a long-term medical application it is an additional bonus if the system does not rely on cables that connect the implant with external components outside the body. Such connections would allow bacteria to enter the skull along the cables and impose a serious infection risk for the patient. Thus power supply and communication between all external and internal components need to be wireless. The best way to supply the internal components with power is not determined yet (e.g. [82, 102, 103, 104, 105]). For the wireless transfer of data two different approaches can be taken: 1.) Data transfer via radio-frequency transmissions (e.g. [82, 102, 106, 107, 108, 109, 110, 111, 112, 113, 114, 115, 116, 117]). 2.) Data transfer via Infrared(IR)-transmissions: Since the skull and skin allows infra-red light to pass [118], it is possible to use IR technology to transmit data [119, 120, 121, 122]. First tests, performed with the BIAS (Bremen Institute for Applied Beam Technology), showed surprisingly high rates (not shown).

On the road of designing such an implant for the long-term goal of realizing a visual cortex prosthesis for blind patients, we envisage that the proposed, external close-loop system will provide a valuable test-bed for flexibly investigating different putative approaches, and for finding suitable specifications for production systems.

### Materials and Methods

For designing the printed circuit board layouts we used Cadsoft Eagle 5.11. For writing the firmware for Orange Tree Technology ZestET1’s FPGA we used Xilinx ISE 14.3 (webpack) and for the Microsemi nano FPGA we applied the Microsemi Libero software (version 11.4). For the simulation of circuit designs we used Texas Instruments TINA v9.3. For creating the 3D rendering of the boards, which were based on the Eagle files, a set of different software tools were used for visualization. For the smaller boards we used Target 3001 V18 and then exported it directly into the pov-file format. For the larger boards we used a combination of Target 3001 V18, the PCB-Pool online 3D Eagle to Step Converter (pcb-pool.com) and the FreeCAD eagle-file import module. These 3D board models were then combined in FreeCAD V0.16 and then exported into the pov-file format. Based on the pov-files, the POV-Ray v3.7 software rendered the pictures. The models for the Samtec connector were taken from the Samtec website.

## Supplementary

In the supplemental data we present the design files for the firmwares, software and PCB designs as Open Source.

## Acknowledgments

We thank Andreas Kreiter, Dieter Gauck, Sunita Mandon, Serge Strokov, Tobias Tessmann, Andreas Schander and Walter Lang from the University of Bremen for fruitfull discussions. This work was supported in part by Bundesministerium fuer Bildung und Forschung Grant 01 GQ 1106 (Bernstein Award Udo Ernst) as well as the Creative Unit I-See ‘The artifical eye: Chronic wireless interface to the visual cortex’ at the University of Bremen.

## Author contributions

D. R., U.E. and K.P. wrote the paper. D.R. designed the boards, firmwares and software as well as tested the system.

## Conflict of interests

The authors declare no conflict of interest. The founding sponsors had no role in the design of the study; in the collection, analyses, or interpretation of data; in the writing of the manuscript, and in the decision to publish the results.

## References

[1] R. P. Finger, B. Bertram, C. Wolfram, and F. G. Holz, “Blindness and visual impairment in germany,” Dtsch Arztebl Int, vol. 109, no. 27–28, pp. 484–489, 2012.

[2] S. N. Flesher, J. L. Collinger, S. T. Foldes, J. M. Weiss, J. E. Downey, E. C. Tyler-Kabara, S. J. Bensmaia, A. B. Schwartz, M. L. Boninger, and R. A. Gaunt, “Intracortical microstimulation of human somatosensory cortex,” Science Translational Medicine, p. aaf8083, 2016.

[3] S. J. Bensmaia and L. E. Miller, “Restoring sensorimotor function through intracortical interfaces: progress and looming challenges,” Nature Reviews Neuroscience, vol. 15, no. 5, pp. 313–325, 2014.

[4] J. E. O’Doherty, M. A. Lebedev, P. J. Ifft, K. Z. Zhuang, S. Shokur, H. Bleuler, and M. A. Nicolelis, “Active tactile exploration using a brain-machine-brain interface,” Nature, vol. 479, no. 7372, pp. 228–231, 2011.

[5] M. A. Lebedev and M. A. Nicolelis, “Brain-machine interfaces: past, present and future,” TRENDS in Neurosciences, vol. 29, no. 9, pp. 536–546, 2006.

[6] T. Lenarz, H. H. Lim, G. Reuter, J. F. Patrick, and M. Lenarz, “The auditory midbrain implant: a new auditory prosthesis for neural deafness-concept and device description,” Otology & Neurotology, vol. 27, no. 6, pp. 838–843, 2006.

[7] T. Moser, “Optogenetic stimulation of the auditory pathway for research and future prosthetics,” Current opinion in neurobiology, vol. 34, pp. 29–36, 2015.

[8] D. R. Moore and R. V. Shannon, “Beyond cochlear implants: awakening the deafened brain,” Nature neuroscience, vol. 12, no. 6, pp. 686–691, 2009.

[9] F.-G. Zeng, S. Rebscher, W. Harrison, X. Sun, and H. Feng, “Cochlear implants: system design, integration, and evaluation,” IEEE reviews in biomedical engineering, vol. 1, pp. 115–142, 2008.

[10] G. S. Brindley and W. Lewin, “The sensations produced by electrical stimulation of the visual cortex,” The Journal of physiology, vol. 196, no. 2, p. 479, 1968.

[11] P. M. Lewis and J. V. Rosenfeld, “Electrical stimulation of the brain and the development of cortical visual prostheses: An historical perspective,” Brain research, vol. 1630, pp. 208–224, 2016.

[12] E. Zrenner, “Fighting blindness with microelectronics,” Science Translational Medicine, vol. 5, no. 210, pp. 210ps16–210ps16, 2013.

[13] R. A. Normann, E. M. Maynard, P. J. Rousche, and D. J. Warren, “A neural interface for a cortical vision prosthesis,” Vision research, vol. 39, no. 15, pp. 2577–2587, 1999.

[14] P. H. Schiller and E. J. Tehovnik, “Visual prosthesis,” Perception, vol. 37, no. 10, pp. 1529–1559, 2008.

[15] E. Schmidt, M. Bak, F. Hambrecht, C. Kufta, D. O’rourke, and P. Vallabhanath, “Feasibility of a visual prosthesis for the blind based on intracortical micro stimulation of the visual cortex,” Brain, vol. 119, no. 2, pp. 507–522, 1996.

[16] J. Coulombe, M. Sawan, et al., “A highly flexible system for microstimulation of the visual cortex: design and implementation,” IEEE transactions on biomedical circuits and systems, vol. 1, no. 4, pp. 258–269, 2007.

[17] Y. H.-L. Luo and L. da Cruz, “A review and update on the current status of retinal prostheses (bionic eye),” British medical bulletin, vol. 109, no. 1, pp. 31–44, 2014.

[18] A. T. Chuang, C. E. Margo, and P. B. Greenberg, “Retinal implants: a systematic review,” British Journal of Ophthalmology, vol. 98, no. 7, pp. 852–856, 2014.

[19] G. Dagnelie, “Retinal implants: emergence of a multidisciplinary field.,” Current opinion in neurology, vol. 25, no. 1, pp. 67–75, 2012.

[20] E. Zrenner, “Artificial vision: solar cells for the blind,” Nature Photonics, vol. 6, no. 6, pp. 344–345, 2012.

[21] J. D. Dorn, A. K. Ahuja, A. Caspi, L. da Cruz, G. Dagnelie, J.-A. Sahel, R. J. Greenberg, M. J. McMahon, A. I. S. Group, et al., “The detection of motion by blind subjects with the epiretinal 60-electrode (argus ii) retinal prosthesis,” JAMA ophthalmology, vol. 131, no. 2, pp. 183–189, 2013.

[22] A. Caspi, J. D. Dorn, K. H. McClure, M. S. Humayun, R. J. Greenberg, and M. J. McMahon, “Feasibility study of a retinal prosthesis: spatial vision with a 16-electrode implant,” Archives of Ophthalmology, vol. 127, no. 4, pp. 398–401, 2009.

[23] J. F. Rizzo III, “Update on retinal prosthetic research: the boston retinal implant project,” Journal of Neuro-ophthalmology, vol. 31, no. 2, pp. 160–168, 2011.

[24] E. Zrenner, R. Wilke, K. Bartz-Schmidt, H. Benav, D. Besch, F. Gekeler, J. Koch, K. Porubska, H. Sachs, and B. Wilhelm, “Blind retinitis pigmentosa patients can read letters and recognize the direction of fine stripe patterns with subretinal electronic implants,” Investigative Ophthalmology & Visual Science, vol. 50, no. 13, pp. 4581–4581, 2009.

[25] K. Stingl, K. U. Bartz-Schmidt, D. Besch, A. Braun, A. Bruckmann, F. Gekeler, U. Greppmaier, S. Hipp, G. Hörtdörfer, C. Kernstock, et al., “Artificial vision with wirelessly powered subretinal electronic implant alpha-ims,” in Proc. R. Soc. B, vol. 280, p. 20130077, The Royal Society, 2013.

[26] J. D. Weiland and M. S. Humayun, “Visual prosthesis,” Proceedings of the IEEE, vol. 96, no. 7, pp. 1076–1084, 2008.

[27] W. H. Dobelle, M. Mladejovsky, and J. Girvin, “Artificial vision for the blind: electrical stimulation of visual cortex offers hope for a functional prosthesis,” Science, vol. 183, no. 4123, pp. 440–444, 1974.

[28] P. M. Lewis, H. M. Ackland, A. J. Lowery, and J. V. Rosenfeld, “Restoration of vision in blind individuals using bionic devices: a review with a focus on cortical visual prostheses,” Brain research, vol. 1595, pp. 51–73, 2015.

[29] N. Cicmil and K. Krug, “Playing the electric light orchestra—how electrical stimulation of visual cortex elucidates the neural basis of perception,” Phil. Trans. R. Soc. B, vol. 370, no. 1677, p. 20140206, 2015.

[30] W. H. Dobelle, “Artificial vision for the blind by connecting a television camera to the visual cortex,” ASAIO journal, vol. 46, no. 1, pp. 3–9, 2000.

[31] K. L. Clark, K. M. Armstrong, and T. Moore, “Probing neural circuitry and function with electrical microstimulation,” Proceedings of the Royal Society of London B: Biological Sciences, p. rspb20102211, 2011.

[32] E. Noorsal, K. Sooksood, H. Xu, R. Hornig, J. Becker, and M. Ortmanns, “A neural stimulator frontend with high-voltage compliance and programmable pulse shape for epiretinal implants,” IEEE Journal of Solid-State Circuits, vol. 47, no. 1, pp. 244–256, 2012.

[33] D. Osipov, S. Paul, S. Strokov, and A. K. Kreiter, “A new current stimulator architecture for visual cortex stimulation,” in Nordic Circuits and Systems Conference (NORCAS): NORCHIP & International Symposium on System-on-Chip (SoC), 2015, pp. 1–4, IEEE, 2015.

[34] A. M. Aravanis, L.-P. Wang, F. Zhang, L. A. Meltzer, M. Z. Mogri, M. B. Schneider, and K. Deisseroth, “An optical neural interface: in vivo control of rodent motor cortex with integrated fiberoptic and optogenetic technology,” Journal of neural engineering, vol. 4, no. 3, p. S143, 2007.

[35] X. Navarro, T. B. Krueger, N. Lago, S. Micera, T. Stieglitz, and P. Dario, “A critical review of interfaces with the peripheral nervous system for the control of neuroprostheses and hybrid bionic systems,” Journal of the Peripheral Nervous System, vol. 10, no. 3, pp. 229–258, 2005.

[36] C. Wang, E. Brunton, S. Haghgooie, K. Cassells, A. Lowery, and R. Rajan, “Characteristics of electrode impedance and stimulation efficacy of a chronic cortical implant using novel annulus electrodes in rat motor cortex,” Journal of neural engineering, vol. 10, no. 4, p. 046010, 2013.

[37] E. K. Brunton, R. Rajan, and A. J. Lowery, “Optimising electrode surface area to minimize power consumption in a cortical penetrating prosthesis,” in Neural Engineering (NER), 2013 6th International IEEE/EMBS Conference on, pp. 1477–1480, IEEE, 2013.

[38] E. K. Brunton, A. J. Lowery, and R. Rajan, “A comparison of microelectrodes for a visual cortical prosthesis using finite element analysis,” Frontiers in Neuroengineering, vol. 5, p. 23, 2012.

[39] R. Samba, T. Herrmann, and G. Zeck, “Pedot–cnt coated electrodes stimulate retinal neurons at low voltage amplitudes and low charge densities,” Journal of neural engineering, vol. 12, no. 1, p. 016014, 2015.

[40] A. Schander, H. Stemmann, E. Tolstosheeva, R. Roese, V. Biefeld, L. Kempen, A. Kreiter, and W. Lang, “Design and fabrication of novel multi-channel floating neural probes for intracortical chronic recording,” Sensors and Actuators A: Physical, 2016.

[41] E. Tolstosheeva, V. Gordillo-González, T. Hertzberg, L. Kempen, I. Michels, A. Kreiter, and W. Lang, “A novel flex-rigid and soft-release ecog array,” in 2011 Annual International Conference of the IEEE Engineering in Medicine and Biology Society, pp. 2973–2976, IEEE, 2011.

[42] A. Schander, T. Teßmann, S. Strokov, H. Stemmann, A. Kreiter, and W. Lang, “In-vitro evaluation of the long-term stability of pedot: Pss coated microelectrodes for chronic recording and electrical stimulation of neurons,” in Engineering in Medicine and Biology Society (EMBC), 2016 IEEE 38th Annual International Conference of the, pp. 6174–6177, IEEE, 2016.

[43] B. Rubehn, C. Bosman, R. Oostenveld, P. Fries, and T. Stieglitz, “A mems-based flexible multichannel ecog-electrode array,” Journal of neural engineering, vol. 6, no. 3, p. 036003, 2009.

[44] E. D. Cohen, “Prosthetic interfaces with the visual system: biological issues,” Journal of neural engineering, vol. 4, no. 2, p. R14, 2007.

[45] N. K. Logothetis, M. Augath, Y. Murayama, A. Rauch, F. Sultan, J. Goense, A. Oeltermann, and H. Merkle, “The effects of electrical microstimulation on cortical signal propagation,” Nature neuroscience, vol. 13, no. 10, pp. 1283–1291, 2010.

[46] E. Tehovnik, A. Tolias, F. Sultan, W. Slocum, and N. Logothetis, “Direct and indirect activation of cortical neurons by electrical microstimulation,” Journal of neurophysiology, vol. 96, no. 2, pp. 512–521, 2006.

[47] E. J. Tehovnik and W. M. Slocum, “Electrical induction of vision,” Neuroscience & Biobehavioral Reviews, vol. 37, no. 5, pp. 803–818, 2013.

[48] D. Brugger, S. Butovas, M. Bogdan, and C. Schwarz, “Real-time adaptive microstimulation increases reliability of electrically evoked cortical potentials,” IEEE Transactions on Biomedical Engineering, vol. 58, no. 5, pp. 1483–1491, 2011.

[49] E. J. Tehovnik and W. M. Slocum, “Phosphene induction by microstimulation of macaque v1,” Brain research reviews, vol. 53, no. 2, pp. 337–343, 2007.

[50] M. H. Histed, A. M. Ni, and J. H. Maunsell, “Insights into cortical mechanisms of behavior from microstimulation experiments,” Progress in neurobioloqy, vol. 103, pp. 115–130, 2013.

[51] J. R. Bartlett, E. A. DeYoe, R. W. Doty, B. B. Lee, J. D. Lewine, N. Negrao, and W. H. Overman, “Psychophysics of electrical stimulation of striate cortex in macaques,” Journal of neurophysiology, vol. 94, no. 5, pp. 3430–3442, 2005.

[52] E. A. DeYoe, J. D. Lewine, and R. W. Doty, “Laminar variation in threshold for detection of electrical excitation of striate cortex by macaques,” Journal of neurophysiology, vol. 94, no. 5, pp. 3443–3450, 2005.

[53] S. Butovas and C. Schwarz, “Spatiotemporal effects of microstimulation in rat neocortex: a parametric study using multielectrode recordings,” Journal of neurophysiology, vol. 90, no. 5, pp. 3024–3039, 2003.

[54] S. Borchers, M. Himmelbach, N. Logothetis, and H.-O. Karnath, “Direct electrical stimulation of human cortex—the gold standard for mapping brain functions?,” Nature Reviews Neuroscience, vol. 13, no. 1, pp. 63–70, 2012.

[55] S. Butovas, S. G. Hormuzdi, H. Monyer, and C. Schwarz, “Effects of electrically coupled inhibitory networks on local neuronal responses to intracortical microstimulation,” Journal of neurophysiology, vol. 96, no. 3, pp. 1227–1236, 2006.

[56] M. Semprini, L. Bennicelli, and A. Vato, “A parametric study of intracortical microstimulation in behaving rats for the development of artificial sensory channels,” in 2012 Annual International Conference of the IEEE Engineering in Medicine and Biology Society, pp. 799–802, IEEE, 2012.

[57] S. Venkatraman, K. Elkabany, J. D. Long, Y. Yao, and J. M. Carmena, “A system for neural recording and closed-loop intracortical microstimulation in awake rodents,” IEEE Transactions on Biomedical Engineering, vol. 56, no. 1, pp. 15–22, 2009.

[58] C. A. Anastassiou, S. M. Montgomery, M. Barahona, G. Buzsáki, and C. Koch, “The effect of spatially inhomogeneous extracellular electric fields on neurons,” The Journal of neuroscience, vol. 30, no. 5, pp. 1925–1936, 2010.

[59] C. A. Anastassiou, R. Perin, H. Markram, and C. Koch, “Ephaptic coupling of cortical neurons,” Nature neuroscience, vol. 14, no. 2, pp. 217–223, 2011.

[60] F. Fröhlich and D. A. McCormick, “Endogenous electric fields may guide neocortical network activity,” Neuron, vol. 67, no. 1, pp. 129–143, 2010.

[61] A. L. Orsborn, H. G. Moorman, S. A. Overduin, M. M. Shanechi, D. F. Dimitrov, and J. M. Carmena, “Closed-loop decoder adaptation shapes neural plasticity for skillful neuroprosthetic control,” Neuron, vol. 82, no. 6, pp. 1380–1393, 2014.

[62] S. Stanslaski, P. Cong, D. Carlson, W. Santa, R. Jensen, G. Molnar, W. J. Marks, A. Shafquat, and T. Denison, “An implantable bi-directional brain-machine interface system for chronic neuroprosthesis research,” in 2009 Annual International Conference of the IEEE Engineering in Medicine and Biology Society, pp. 5494–5497, IEEE, 2009.

[63] R. R. Harrison and C. Charles, “A low-power low-noise cmos amplifier for neural recording applications,” IEEE Journal of solid-state circuits, vol. 38, no. 6, pp. 958–965, 2003.

[64] F. Zhang, J. Holleman, and B. P. Otis, “Design of ultra-low power biopotential amplifiers for biosignal acquisition applications,” IEEE transactions on biomedical circuits and systems, vol. 6, no. 4, pp. 344–355, 2012.

[65] X. Zhang, W. Pei, B. Huang, J. Chen, N. Guan, and H. Chen, “Implantable microsystem for wireless neural recording applications,” in Complex Medical Engineering, 2009. CME. ICME International Conference on, pp. 1–4, IEEE, 2009.

[66] R. R. Harrison, “The design of integrated circuits to observe brain activity,” Proceedings of the IEEE, vol. 96, no. 7, pp. 1203–1216, 2008.

[67] W. Wattanapanitch and R. Sarpeshkar, “A low-power 32-channel digitally programmable neural recording integrated circuit,” IEEE Transactions on Biomedical Circuits and Systems, vol. 5, no. 6, pp. 592–602, 2011.

[68] F. Shahrokhi, K. Abdelhalim, D. Serletis, P. L. Carlen, and R. Genov, “The 128-channel fully differential digital integrated neural recording and stimulation interface,” IEEE Transactions on Biomedical Circuits and Systems, vol. 4, no. 3, pp. 149–161, 2010.

[69] C. M. Lopez, D. Prodanov, D. Braeken, I. Gligorijevic, W. Eberle, C. Bartic, R. Puers, and G. Gielen, “A multichannel integrated circuit for electrical recording of neural activity, with independent channel programmability,” IEEE transactions on biomedical circuits and systems, vol. 6, no. 2, pp. 101–110, 2012.

[70] F. Zhang, M. Aghagolzadeh, and K. Oweiss, “A low-power implantable neuroprocessor on nano-fpga for brain machine interface applications,” in 2011 IEEE International Conference on Acoustics, Speech and Signal Processing (ICASSP), pp. 1593–1596, IEEE, 2011.

[71] M. J. Kane, P. P. Breen, F. Quondamatteo, and G. OLaighin, “Bion microstimulators: A case study in the engineering of an electronic implantable medical device,” Medical engineering & physics, vol. 33, no. 1, pp. 7–16, 2011.

[72] B. Gosselin, “Recent advances in neural recording microsystems,” Sensors, vol. 11, no. 5, pp. 4572–4597, 2011.

[73] K. D. Wise, “Wireless integrated microsystems: Wearable and implantable devices for improved health care,” in TRANSDUCERS 2009–2009 International Solid-State Sensors, Actuators and Microsystems Conference, pp. 1–8, IEEE, 2009.

[74] V. Gilja, C. A. Chestek, I. Diester, J. M. Henderson, K. Deisseroth, and K. V. Shenoy, “Challenges and opportunities for next-generation intracortically based neural prostheses,” IEEE Transactions on Biomedical Engineering, vol. 58, no. 7, pp. 1891–1899, 2011.

[75] F. J. Lane, K. B. Nitsch, and P. Troyk, “Participant perspectives from a cortical vision implant study: Ethical and psychological implications,” in 2015 7th International IEEE/EMBS Conference on Neural Engineering (NER), pp. 264–267, IEEE, 2015.

[76] A. J. Lowery, J. V. Rosenfeld, P. M. Lewis, D. Browne, A. Mohan, E. Brunton, E. Yan, J. Maller, C. Mann, R. Rajan, et al., “Restoration of vision using wireless cortical implants: The monash vision group project,” in 2015 37th Annual International Conference of the IEEE Engineering in Medicine and Biology Society (EMBC), pp. 1041–1044, IEEE, 2015.

[77] S. Strokov, H. Stemmann, D. Osipov, S. Paul, T. Tessmann, A. Schander, W. Lang, and A. Kreiter, “Switchable low-cost multichannel neural stimulation module for parallel recording and stimulation doi: 10.13140/rg.2.2.14422.86081,” in 11. Bernstein Sparks Workshop: Naturalistic integration of information from external stimulation into the ongoing neuronal activities of the brain, 2016.

[78] D. McCreery, W. Agnew, T. Yuen, and L. Bullara, “Comparison of neural damage induced by electrical stimulation with faradaic and capacitor electrodes,” Annals of biomedical engineering, vol. 16, no. 5, pp. 463–481, 1988.

[79] TI, “Texas instruments: A comprehensive study of the howland current,” Application Report, vol. SNOA474A, 2008.

[80] IPC, “Ipc-2221a generic standard on printed board design,” Association Connecting Electronics Industries, 2003.

[81] D. Osipov and S. Paul, “A novel hv-switch scheme with gate-source overvoltage protection for bidirectional neural interfaces,” in 2015 IEEE International Conference on Electronics, Circuits, and Systems (ICECS), pp. 25–28, IEEE, 2015.

[82] D. Rotermund, J. Pistor, J. Hoeffmann, T. Schellenberg, D. Boll, E. Tolstosheeva, D. Gauck, D. Peters-Drolshagen, A. K. Kreiter, M. Schneider, et al., “Open hardware: Towards a fully-wireless sub-cranial neuro-implant for measuring electrocorticography signals,” bioRxiv, p. 036855, 2016.

[83] M. Yin, D. A. Borton, J. Aceros, W. R. Patterson, and A. V. Nurmikko, “A 100-channel hermetically sealed implantable device for chronic wireless neurosensing applications,” IEEE transactions on biomedical circuits and systems, vol. 7, no. 2, pp. 115–128, 2013.

[84] D. A. Borton, M. Yin, J. Aceros, and A. Nurmikko, “An implantable wireless neural interface for recording cortical circuit dynamics in moving primates,” Journal of neural engineering, vol. 10, no. 2, p. 026010, 2013.

[85] M. Hirata, K. Matsushita, T. Suzuki, T. Yoshida, S. Morris, T. Yanagisawa, M. Kawato, and T. Yoshimine, “A fully-implantable wireless system for human brain–machine interfaces using brain surface electrodes: W-herbs,” IEICE transactions on communications, vol. 94, no. 9, pp. 2448–2453, 2011.

[86] A. Sharma, L. Rieth, P. Tathireddy, R. Harrison, H. Oppermann, M. Klein, M. Töpper, E. Jung, R. Normann, G. Clark, et al., “Evaluation of the packaging and encapsulation reliability in fully integrated, fully wireless 100 channel utah slant electrode array (usea): Implications for long term functionality,” Sensors and Actuators A: Physical, vol. 188, pp. 167–172, 2012.

[87] R. R. Harrison, R. J. Kier, C. A. Chestek, V. Gilja, P. Nuyujukian, S. Ryu, B. Greger, F. Solzbacher, and K. V. Shenoy, “Wireless neural recording with single low-power integrated circuit,” IEEE transactions on neural systems and rehabilitation engineering, vol. 17, no. 4, pp. 322–329, 2009.

[88] R. R. Harrison, R. J. Kier, S. Kim, L. Rieth, D. J. Warren, N. M. Ledbetter, G. A. Clark, F. Solzbacher, C. A. Chestek, V. Gilja, et al., “A wireless neural interface for chronic recording,” in 2008 IEEE Biomedical Circuits and Systems Conference, pp. 125–128, IEEE, 2008.

[89] S. Kim, R. Bhandari, M. Klein, S. Negi, L. Rieth, P. Tathireddy, M. Toepper, H. Oppermann, and F. Solzbacher, “Integrated wireless neural interface based on the utah electrode array,” Biomedical microdevices, vol. 11, no. 2, pp. 453–466, 2009.

[90] E. Okada and D. T. Delpy, “Near-infrared light propagation in an adult head model. ii. effect of superficial tissue thickness on the sensitivity of the near-infrared spectroscopy signal,” Applied optics, vol. 42, no. 16, pp. 2915–2921, 2003.

[91] A. Tamura, S. Hayashi, K. Nagayama, and T. Matsumoto, “Mechanical characterization of brain tissue in high-rate extension,” Journal of Biomechanical Science and Engineering, vol. 3, no. 2, pp. 263–274, 2008.

[92] K. Miller and K. Chinzei, “Mechanical properties of brain tissue in tension,” Journal of biomechanics, vol. 35, no. 4, pp. 483–490, 2002.

[93] K. Miller, K. Chinzei, G. Orssengo, and P. Bednarz, “Mechanical properties of brain tissue in-vivo: experiment and computer simulation,” Journal of biomechanics, vol. 33, no. 11, pp. 1369–1376, 2000.

[94] K. M. Silay, C. Dehollain, and M. Declercq, “Numerical analysis of temperature elevation in the head due to power dissipation in a cortical implant,” in 2008 30th Annual International Conference of the IEEE Engineering in Medicine and Biology Society, pp. 951–956, IEEE, 2008.

[95] S. Kim, P. Tathireddy, R. A. Normann, and F. Solzbacher, “In vitro and in vivo study of temperature increases in the brain due to a neural implant,” in 2007 3rd International IEEE/EMBS Conference on Neural Engineering, pp. 163–166, IEEE, 2007.

[96] S. Kim, P. Tathireddy, R. A. Normann, and F. Solzbacher, “Thermal impact of an active 3-d microelectrode array implanted in the brain,” IEEE Transactions on Neural Systems and Rehabilitation Engineering, vol. 15, no. 4, pp. 493–501, 2007.

[97] V. S. Polikov, P. A. Tresco, and W. M. Reichert, “Response of brain tissue to chronically implanted neural electrodes,” Journal of neuroscience methods, vol. 148, no. 1, pp. 1–18, 2005.

[98] G. Jiang and D. D. Zhou, “Technology advances and challenges in hermetic packaging for implantable medical devices,” in Implantable Neural Prostheses 2, pp. 27–61, Springer, 2009.

[99] W.-s. Lee, J.-K. Lee, S.-A. Lee, J.-K. Kang, and T.-s. Ko, “Complications and results of subdural grid electrode implantation in epilepsy surgery,” Surgical neurology, vol. 54, no. 5, pp. 346–351, 2000.

[100] D. R. Nair, R. Burgess, C. C. McIntyre, and H. Lüders, “Chronic subdural electrodes in the management of epilepsy,” Clinical neurophysiology, vol. 119, no. 1, pp. 11–28, 2008.

[101] J. Voges, Y. Waerzeggers, M. Maarouf, R. Lehrke, A. Koulousakis, D. Lenartz, and V. Sturm, “Deep-brain stimulation: long-term analysis of complications caused by hardware and surgery—experiences from a single centre,” Journal of Neurology, Neurosurgery & Psychiatry, vol. 77, no. 7, pp. 868–872, 2006.

[102] J. Parramon i Piella and E. Valderrama Valles, Energy management, wireless and system solutions for highly integrated implantable devices. Universitat Autònoma de Barcelona,, 2003.

[103] J. Olivo, S. Carrara, and G. De Micheli, “Energy harvesting and remote powering for implantable biosensors,” IEEE Sensors Journal, vol. 11, no. EPFL-ARTICLE-152140, pp. 1573–1586, 2011.

[104] B. I. Rapoport, J. T. Kedzierski, and R. Sarpeshkar, “A glucose fuel cell for implantable brain-machine interfaces,” PloS one, vol. 7, no. 6, p. e38436, 2012.

[105] S. B. Lee, H.-M. Lee, M. Kiani, U.-M. Jow, and M. Ghovanloo, “An inductively powered scalable 32-channel wireless neural recording system-on-a-chip for neuroscience applications,” IEEE transactions on biomedical circuits and systems, vol. 4, no. 6, pp. 360–371, 2010.

[106] J. E. Ferguson and A. D. Redish, “Wireless communication with implanted medical devices using the conductive properties of the body,” Expert review of medical devices, vol. 8, no. 4, pp. 427–433, 2011.

[107] F. Asgarian and A. M. Sodagar, Wireless telemetry for implantable biomedical microsystems. INTECH Open Access Publisher, 2011.

[108] M. L. Navaii, M. Jalali, and H. Sadjedi, “An ultra-low power rf interface for wireless-implantable microsystems,” Microelectronics Journal, vol. 43, no. 11, pp. 848–856, 2012.

[109] M. A. Hannan, S. M. Abbas, S. A. Samad, and A. Hussain, “Modulation techniques for biomedical implanted devices and their challenges,” Sensors, vol. 12, no. 1, pp. 297–319, 2011.

[110] Y.-H. Liu, C.-L. Li, and T.-H. Lin, “A 200-pj/b mux-based rf transmitter for implantable multichannel neural recording,” IEEE Transactions on Microwave Theory and Techniques, vol. 57, no. 10, pp. 2533–2541, 2009.

[111] H. N. Schwerdt, W. Xu, S. Shekhar, A. Abbaspour-Tamijani, B. C. Towe, F. A. Miranda, and J. Chae, “A fully passive wireless microsystem for recording of neuropotentials using rf backscattering methods,” Journal of Microelectromechanical Systems, vol. 20, no. 5, pp. 1119–1130, 2011.

[112] Y.-K. Song, D. A. Borton, S. Park, W. R. Patterson, C. W. Bull, F. Laiwalla, J. Mislow, J. D. Simeral, J. P. Donoghue, and A. V. Nurmikko, “Active microelectronic neurosensor arrays for implantable brain communication interfaces,” IEEE transactions on neural systems and rehabilitation engineering: a publication of the IEEE Engineering in Medicine and Biology Society, vol. 17, no. 4, p. 339, 2009.

[113] H. Miranda, V. Gilja, C. A. Chestek, K. V. Shenoy, and T. H. Meng, “Hermesd: A high-rate long-range wireless transmission system for simultaneous multichannel neural recording applications,” IEEE Transactions on Biomedical Circuits and Systems, vol. 4, no. 3, pp. 181–191, 2010.

[114] R. R. Harrison, H. Fotowat, R. Chan, R. J. Kier, R. Olberg, A. Leonardo, and F. Gabbiani, “Wireless neural/emg telemetry systems for small freely moving animals,” IEEE transactions on biomedical circuits and systems, vol. 5, no. 2, pp. 103–111, 2011.

[115] S. J. Thomas, R. R. Harrison, A. Leonardo, and M. S. Reynolds, “A battery-free multichannel digital neural/emg telemetry system for flying insects,” IEEE transactions on biomedical circuits and systems, vol. 6, no. 5, pp. 424–436, 2012.

[116] H. Sato, C. W. Berry, Y. Peeri, E. Baghoomian, B. E. Casey, G. Lavella, J. M. VandenBrooks, J. Harrison, and M. M. Maharbiz, “Remote radio control of insect flight,” Frontiers in integrative neuroscience, vol. 3, p. 24, 2009.

[117] M. S. Chae, Z. Yang, M. R. Yuce, L. Hoang, and W. Liu, “A 128-channel 6 mw wireless neural recording ic with spike feature extraction and uwb transmitter,” IEEE Transactions on Neural Systems and Rehabilitation Engineering, vol. 17, no. 4, pp. 312–321, 2009.

[118] M. Firbank, M. Hiraoka, M. Essenpreis, and D. Delpy, “Measurement of the optical properties of the skull in the wavelength range 650–950 nm,” Physics in medicine and biology, vol. 38, no. 4, p. 503, 1993.

[119] J. Aceros, M. Yin, D. A. Borton, W. R. Patterson, and A. V. Nurmikko, “A 32-channel fully implantable wireless neurosensor for simultaneous recording from two cortical regions,” in 2011 Annual International Conference of the IEEE Engineering in Medicine and Biology Society, pp. 2300–2306, IEEE, 2011.

[120] Y. Damestani, C. L. Reynolds, J. Szu, M. S. Hsu, Y. Kodera, D. K. Binder, B. H. Park, J. E. Garay, M. P. Rao, and G. Aguilar, “Transparent nanocrystalline yttria-stabilized-zirconia calvarium prosthesis,” Nanomedicine: Nanotechnology, Biology and Medicine, vol. 9, no. 8, pp. 1135–1138, 2013.

[121] D. M. Ackermann Jr, High speed transcutaneous optical telemetry link. PhD thesis, Case Western Reserve University, 2008.

[122] K.-W. Cheng, X. Zou, J. H. Cheong, R.-F. Xue, Z. Chen, L. Yao, H.-K. Cha, S. J. Cheng, P. Li, L. Liu, et al., “100-channel wireless neural recording system with 54mb/s data link and 40%-efficiency power link,” in Solid State Circuits Conference (A-SSCC), 2012 IEEE Asian, pp. 185–188, IEEE, 2012.

